# Genetic regulation of human aortic smooth muscle cell gene expression and splicing predict causal coronary artery disease genes

**DOI:** 10.1101/2022.01.24.477536

**Authors:** Rédouane Aherrahrou, Dillon Lue, R Noah Perry, Yonathan Tamrat Aberra, Mohammad Daud Khan, Joon Yuhl Soh, Tiit Örd, Prosanta Singha, Huda Gilani, Ernest Diez Benavente, Doris Wong, Jameson Hinkle, Lijiang Ma, Gloria M Sheynkman, Hester M den Ruijter, Clint L Miller, Johan LM Björkegren, Minna U Kaikkonen, Mete Civelek

## Abstract

Coronary artery disease (CAD) is the leading cause of death worldwide. Recent meta-analyses of genome-wide association studies (GWAS) have identified over 175 loci associated with CAD. The majority of these loci are in non-coding regions and are predicted to regulate gene expression. Given that vascular smooth muscle cells (SMCs) play critical roles in the development and progression of CAD, we hypothesized that a subset of the CAD GWAS risk loci are associated with the regulation of transcription in distinct SMC phenotypes. Here, we measured gene expression in SMCs isolated from the ascending aortas of 151 ethnically diverse heart transplant donors in quiescent or proliferative conditions and calculated the association of their expression and splicing with ∼6.3 million imputed single nucleotide polymorphism (SNP) markers across the genome. We identified 4,910 expression and 4,412 splice quantitative trait loci (sQTL) that represent regions of the genome associated with transcript abundance and splicing. 3,660 of the eQTLs had not been observed in the publicly available Genotype-Tissue Expression dataset. Further, 29 and 880 of the eQTLs were SMC- and sex-specific, respectively. To identify the effector transcript(s) regulated by CAD GWAS loci, we used four distinct colocalization approaches and identified 84 eQTL and 164 sQTLs that colocalized with CAD loci, highlighting the importance of genetic regulation of mRNA splicing as a molecular mechanism for CAD genetic risk. Notably, 20% and 35% of the eQTLs were unique to quiescent or proliferative SMCs, respectively. Two CAD loci colocalized with a SMC sex-specific eQTL (*AL160313.1* and *TERF2IP*) and another locus colocalized with SMC-specific eQTL (*ALKBH8*). Also, 27% and 37% of the sQTLs were unique to quiescent or proliferative SMCs, respectively. The most significantly associated CAD locus, 9p21, was an sQTL for the long non-coding RNA *CDKN2B-AS1*, also known as *ANRIL*, in proliferative SMCs. Collectively, these results provide evidence for the molecular mechanisms of genetic susceptibility to CAD in distinct SMC phenotypes.

## INTRODUCTION

Coronary artery disease (CAD) is the leading cause of death worldwide^1^. Heritability estimates for CAD vary between 40% to 70%, suggesting a strong genetic contribution to disease pathology^2^. Genome-wide association studies (GWAS) have identified 175 loci associated with increased risk for CAD^3–5^. Approximately 40% of the CAD loci are associated with known risk factors, such as blood lipid levels, nitric oxide signaling, and blood pressure^3^. The remaining 60% have unknown mechanisms, but there is evidence that some of these loci function through the vessel wall where the disease develops^6^. In addition, 94% of CAD-associated genetic variants are in non-coding regions of the genome^7^, implying that the disease-causing loci may involve regulatory mechanisms that affect the transcription of genes^6^. Therefore, gene expression studies performed in cells and tissues relevant to CAD in human populations can pinpoint the regulatory mechanisms of disease susceptibility^8^.

Expression quantitative trait loci (eQTL) and splicing quantitative trait loci (sQTL) analyses are key approaches that link genetic variants with variations in gene expression and splicing patterns, respectively^8,9^. They have enabled the prioritization of genetic variants within GWAS loci for different traits, and have shown that trait-associated genetic variants often function in a tissue-or cell type-specific manner^10–13^. Smooth muscle cells (SMCs), which make up the medial layer of arteries, play key roles in the integrity of the vessel wall, regulation of blood pressure and initiation and development of atherosclerosis. Recent studies provided compelling evidence that SMCs can play either beneficial or detrimental roles in lesion pathogenesis depending on the nature of their phenotypic changes^14–16^. In addition, SMC phenotypic switching seems to be important in explaining sex differences in atherosclerotic plaque composition^17^. Thus, identifying the genetic determinants of SMC gene expression is crucial for understanding the biological significance of CAD-associated genetic variants functioning in SMCs. This will also inform the prediction of novel drug candidates targeting the disease processes in the vessel wall.

Despite publicly available population-level gene expression datasets from various tissues and cells, including 54 tissues from the Genotype-Tissue Expression (GTEx) project^18^ and atherosclerosis-relevant tissues and cell type from the STARNET cohort^19^, aortic endothelial cells^20,21^, monocytes^22^, whole blood^23^, and coronary artery SMCs^24^ as well as epigenome profiling from the Roadmap Epigenomics Project^25^, more than half of the CAD loci are still not functionally annotated. Therefore, in this study, using a multi-omics approach, we identified the genetic variants that are associated with SMC-specific gene expression derived from 151 healthy and ethnically-diverse heart transplant donors. This allowed us to identify CAD-associated loci, including the most significantly associated CAD locus, 9p21, that perturb SMC transcription, thereby leading to the prediction of candidate effector transcripts and candidate causal variants in these disease-associated loci.

## MATERIALS and METHODS

### Cell culture

We isolated SMCs enzymatically from the explants of ascending aortas from 136 healthy heart transplant donors (118 males and 33 females) at the University of California at Los Angeles (UCLA) transplant program as described previously^26^. We also purchased aortic SMCs isolated from ascending aortas from 15 donors from Lonza and PromoCell. We maintained the cells in Smooth Muscle Cell Basal Medium (SmBM, CC-3181, Lonza) supplemented with Smooth Muscle Medium-2 SingleQuots Kit (SmGM-2, CC-4149, Lonza) (complete media). We cultured the SMCs in complete media (containing 5% FBS) until 90% confluence. We then switched to either serum-free media to mimic the quiescent state of SMCs in healthy arteries or continued to culture in complete media to mimic the proliferative state of SMCs in atherosclerotic arteries for 24 hours^27,28^. We harvested the cells in Qiazol and extracted total RNA. The Institutional Review Boards of UCLA and the University of Virginia approved this study.

### Gene silencing and proliferation

We transfected human coronary artery smooth muscle cells (HCASMC), which were maintained in M231 medium (Gibco, M231500) supplemented with SMGS (Gibco, S00725), with control (ThermoFisher, 4390846) or SNHG18 (s452763) siRNA using oligofectamine (Invitrogen, 12252011) per manufacturer’s standard protocol. To measure the extent of downregulation, we collected the cells after 48 hours and extracted the RNA using QIAGEN RNeasy Plus Mini Kit and performed qPCR for GAPDH and SNHG18. We used the following primer pairs: GAPDH-F: TCGGAGTCAACGGATTTG and GAPDH-R: CAACAATATCCACTTTACCAGAG; SNHG18-F: ATGACTGTGGGCCATGAGTG and SNHG18-R: AAAGCAGCCCTAGGCAATCT. ΔΔCt method was used to calculate the relative gene expression of SNHG18 compared to GAPDH housekeeping gene. We performed the proliferation assay in a 96-well plate using the Incucyte S3 Live-Cell Analysis System with the Incucyte Nuclight Rapid Red Dye for nuclear labeling. To monitor cell proliferation, we performed imaging every two hours for four days.

### Genotyping and ancestry determination

We genotyped the donors using the Illumina Multi-Ethnic Global genotyping array for 1.8 million single nucleotide polymorphisms (SNPs). We pruned the SNPs based on call rate (< 2%), Hardy-Weinberg equilibrium (P_HWE_<1×10^−6^), and minor allele frequency (< 5%), and imputed non-genotyped SNPs using the 1000 Genomes Phase 3 reference panel^29^. After removing the SNPs with minor allele frequency less than 5% and imputation quality less than 0.3, we were left with ∼6.3 million SNPs for association studies. To determine the ancestral background of the donors, we excluded SNPs in regions of extended high linkage disequilibrium (LD) and pruned the remaining SNPs at an LD threshold R^2^ > 0.2. We clustered the filtered genotypes with the genotypes of 12 populations represented in the 1000 Genomes Phase 3 data^29^ with principal component analysis (PCA) implemented in KING^30^.

### RNA extraction, sequencing, mapping and quantification

We performed the sequencing of the ribosomal RNA-depleted total RNA isolated from SMCs of the 151 donors cultured in the presence or absence of 5% FBS. Total RNA was extracted using the RNeasy Micro Kit (Qiagen) and the RNase-free DNase Set. RNA integrity scores for all samples, as measured by the Agilent TapeStation, were greater than 9, indicating high-quality RNA preparations. Sequencing libraries were prepared with the Illumina TruSeq Stranded mRNA Library Prep Kit and were sequenced to ∼100 million read depth with 150 bp paired-end reads at the Psomogen sequencing facility. We trimmed the reads with low average Phred scores (<20) using Trim Galore and mapped the reads to the hg38 version of the human reference genome using the STAR Aligner^31^ in two-pass mode to increase the mapping efficiency and sensitivity. We only retained the uniquely mapped read pairs. We quantified gene expression by calculating the transcripts per million (TPM) for each gene using RNA-SeQC^32^ based on GENCODE v32 transcript annotations. In addition to protein-coding RNAs, we also measured the non-coding RNA since they have been shown to play significant roles in SMC biology^33^. We considered a gene as expressed if it had more than 6 read counts and 0.1 TPM in at least 20% of the samples.

### Sample swap identification

To detect sample swaps, we used NGSCheckMate^34^ and verifyBamID^35^ to call variants from RNA-seq data and assign the best matches between the RNA-seq and genotype data. This led to the removal of 11 and 6 samples from quiescent (without FBS) and proliferative (with FBS) SMC cultures, respectively.

### Differential gene expression and functional enrichment analysis

We included 14,341 genes with > 6 reads in at least 80% of the samples in at least one of the two conditions for differential expression analysis using DESeq2^36^. We considered genes to be differentially expressed between proliferative and quiescent conditions when P_adj_< 3×10^−3^ and the absolute value of log_2_(fold-change)> 0.5. We identified the surrogate variables using the *svaseq* function in the *sva* package^37^ using gene expression measured in TPM as input. We performed principal component analysis (PCA) analysis of these surrogate variables using the ARSyNSeq function from the NOISeq package in R^38^. To characterize the functional consequences of gene expression changes associated with proliferative and quiescence conditions, we performed Gene Ontology (GO) enrichment analysis. We ranked the genes based on fold-change and differential expression between the two conditions (P_adj_<0.05) and used Gene Set Enrichment Analysis (GSEA) on the ranked set of genes using all expressed genes in SMCs as background.

### *Cis*-eQTL and -sQTL identification

For *cis*-eQTL discovery, we considered the genes with sufficient expression level (TPM > 0.1 and read count > 6) in at least 20% of the samples. After filtering, we normalized the read counts using the trimmed mean of M values (TMM)^39^ followed by inverse normalization. We corrected the gene expression data for technical artifacts and unknown technical confounders using the probabilistic estimation of expression residuals (PEER) framework^35^. To optimize for *cis*-eQTL discovery, we performed eQTL mapping using inverse normalized gene expression residuals corrected with 5, 10, 15, 20, 25, 30, 35, 40 or 45 PEER factors along with sex and 4 genotype principal components (PCs) using tensorQTL permutation pass analysis^41^. We implemented tensorQTL permutation testing to detect the top nominal associated SNP within 1 MB of the transcription start site of a gene, defined as the *cis* region, and with a beta approximation to model the permutation result and correct for all SNPs in linkage disequilibrium (LD) with the most significant SNP (referred here as eSNP) per gene. We used the ‘--permute 1000 10000’ option in tensorQTL. Beta approximated permutation *P*-values were then corrected for multiple testing using the q-value false discovery rate FDR correction^42^. A gene with a *cis*-eQTL (eGene) was defined by having an FDR q-value <0.05. We report the results of eQTL mapping for 30 and 35 PEER factors for the quiescent and proliferative conditions, respectively since we discovered the maximum number of eQTL genes with that many PEER factors.

We utilized LeafCutter^43^ to obtain and quantify clusters of variably spliced introns and tensorQTL to map sQTLs within a 200 KB window around splice donor sites, controlling for sex, four genotype PCs, and 6 and 8 PEER factors for quiescent and proliferative conditions, respectively. We identified secondary and beyond independent eQTLs and sQTLs by rerunning permutation tests in tensorQTL^44^ for every gene or intron, respectively, conditioning on the primary eSNP. Conditional secondary and beyond molecular QTLs were considered significant if the FDR q-value <0.05. We used LocusZoom for the regional visualization of eQTL/sQTL results on the basis of linkage disequilibrium (LD) ascertained from the 151 donors in our study^45^.

### Detecting condition- and sex-biased eQTLs

To determine *cis*-eQTL SNPs with statistically significant differential effects on gene expression in quiescent or proliferative conditions, we first determined the SNP with the highest statistical significance per each *cis*-eQTL gene in either the quiescent or proliferative condition. We then tested if the effect sizes of the eSNP on the *cis*-eGene were significantly different between the two conditions using a Z-test utilizing the effect size (β) and its standard error (σ^2^) as described previously^46^. We corrected the resulting *P*-values for multiple testing using the q-value false discovery rate FDR correction^42^. We determined a condition-specific eQTL if the FDR q-value in the Z-test was < 0.05. 90% confidence intervals of the effect size (β) were calculated for each condition. We considered an effect as positive if the confidence interval did not include 0 and the z-score was positive and an effect as negative if the confidence interval did not include 0 and the z-score was negative. We classified a condition-specific eQTL as condition-specific direction if the confidence intervals differed in sign, condition-specific magnitude if the confidence intervals were the same sign, and condition-specific effect if one of the confidence intervals included 0.

To determine *cis*-eQTL SNPs with statistically significant different effects on gene expression in males and females, we performed sex-biased *cis*-eQTL analysis on autosomal genes in quiescent and proliferative conditions separately. Like standard eQTL mapping, we first normalized read counts using the trimmed mean of M values and inverse normal transformed gene expression data. Next, we used a linear regression model including genotype, 4 genotype PCs, and the same number of PEER factors we used for standard eQTL mapping using tensorQTL, after removing the effect of sex, to test for significance of genotype-by-sex (G x Sex) interaction on expression. We applied eigenMT, a permutation method, that estimates the effective number of independent tests based on the local LD structure^47^. We considered a sex-biased *cis*-eQTL significant if the eigenMT value <0.05. To classify the identified sex-biased *cis*-eQTLs, we conducted independent linear regression models for each sex-biased eQTL gene and eSNP pair, controlling for four genotype PCs and same number of PEER factors we used for sex-biased *cis*-eQTLs. Similar to condition-specific eQTL analysis, we calculated the 90% confidence intervals of the effect size (β) for each sex per condition. We separated the sex-biased eQTLs into three different categories: sex-biased effect, sex-biased direction, and sex-biased magnitude.

### Overlap of SMC cis-eQTLs with GTEx cis-eQTLs and identification of SMC-specific eQTLs

To compare GTEx eQTLs with our SMC eQTLs we utilized the QTlizer R package^48^ to query the significant SMC eSNPs for eQTL signals in GTEx v8^49^. We only retained GTEx eQTL signals at 5% FDR across all the tissues. We used the variant and gene pairs as identifiers in both GTEx and SMC eQTL datasets. We considered any variant and gene pair significant novel if it was only present in our study. If it was found in any GTEx tissues and our study, we considered it as shared.

Additionally, we identified SMC-specific eQTLs with respect to the GTEx eQTL results^49^. For each SMC gene with an eQTL, we selected the most significantly associated SNP, and performed multi-tissue eQTL calling using all the available SNPs within 1 MB of the TSS of the all the genes in the GTEx dataset using METASOFT^50^. We calculated the posterior probability that the effect exists in each tissue as denoted by the m-value. We defined SMC-specific eQTLs as SNP-gene pairs with m-value>0.9 for SMCs and < 0.1 for all the GTEx tissues. We also queried the SMC-specific eQTLs identified by METASOFT for significant associations in the STARNET dataset^19^.

### Colocalization between molecular SMC QTLs and CAD GWAS signals

We examined whether each *cis*-eQTL and sQTL is colocalized with the GWAS loci associated with CAD using four different methods. First, we calculated the linkage disequilibrium (LD) R^2^ value between the GWAS index variant and the variant with the most significant association with expression or splice QTL (eSNP and sSNP) in our study population. We defined GWAS-coincident eQTLs/sQTLs as loci with pairwise LD R^2^ > 0.8 (1000G EUR) between the GWAS index variant and the lead eSNP/sSNP. To evaluate association between the GWAS variant and the lead eQTL variant at each locus, we performed conditional analyses; we tested the association between the index variant and transcript level when the lead GWAS SNP was included in the model. Second, we used Summary Level Mendelian Randomization (SMR)^51^ to test for the pleiotropic association of gene expression in SMCs and CAD. For the eQTL analysis, we performed a SMR test on genes with a *cis*-eQTL *P*-value < 1.42×10^−5^ and 1.69×10^−5^ in the quiescent and proliferative datasets, for a total of 2,228 and 3,090 tests, respectively. For the sQTL datasets, we used a *P-*value cutoff of 1.73×10^−5^ and 3.08×10^−5^ for a total of 6,787 and 7,945 SMR tests in the quiescent and proliferative sQTL datasets, respectively. We used the 1000 Genomes European reference panel to account for linkage disequilibrium. We considered loci with an adjusted SMR *P*-value (5% FDR) to have evidence of colocalization. Third, we used eQTL and GWAS CAusal Variants Identification in Associated Regions (eCAVIAR)^52^, to test for causal SNPs between our eQTL and sQTL data and the CAD GWAS considering variants within a 500 kb and 200 kb window around the eSNP for each eGene or sGene, respectively. The maximum number of causal variants was set to 2 and variants were considered colocalized if the colocalization posterior probability (CLPP) was greater than 0.01. Finally, we implemented Bayesian Colocalization Analysis using Bayes Factors (COLOC) using the R package COLOC^53^. We first selected SNPs in each CAD GWAS locus with genome-wide significance, and then created 200 KB windows around significant SNPs. Following this, we merged nearby windows (>100 KB distance) together to form loci. We input these loci into COLOC with the default priors (p1/p2 = 1 × 10^−4^, and p12 = 1 × 10^−5^), and considered a locus colocalized if PPH4, the hypothesis of a single shared causal variant for both traits within a window, was greater than 0.50. We then plotted and visually inspected all analyzed loci using LocusCompare^54^. Loci that passed both visual inspection and colocalization criteria were considered colocalized.

### Identification of accessible chromatin regions in SMCs and transcription factor binding site analysis

We performed transposase-accessible chromatin with high-throughput sequencing (ATACseq) in SMCs from five donors cultured in quiescent or proliferative conditions. We isolated 50,000 nuclei and incubated them for 30 minutes with hyperactive Tn5 transposase following the Omni-ATAC protocol^55^. We then amplified the transposed DNA for 8 cycles and inspected for fragment length distribution using an Agilent Bioanalyzer High Sensitivity DNA Chip (Agilent Genomics). This revealed the expected nucleosomal laddering pattern, with subnucleosomal, mononucleosomal, and dinucleosomal fragments enriched at 200, 350, and 550 bp, respectively. We then performed 75 bp paired-end sequencing using Illumina NextSeq 500. We aligned the reads to the human reference genome using Bowtie2^56^, removed the mitochondrial DNA reads, retained the uniquely mapped reads, and removed duplicates with SAMTools^57^. After these quality control steps, >95% of the reads mapped to the human genome. We merged the BAM files of individual samples for each culture condition. We called the accessible chromatin region peaks for each of the two merged files using MACS2^58^ with parameters -f BAMPE and -q 0.1. Fraction of reads in called peak regions were greater than 0.3. We overlapped the genomic coordinates of SNPs of interest with ATACseq peaks using BEDTools intersect^59^.

To evaluate whether eQTL SNPs overlapped with putative transcription factor (TF) binding sites, we overlapped eQTLs located in accessible chromatin regions with sites within TF consensus motifs. We utilized the SNP2TFBS resource of estimated effects of SNPs on predicted TF binding, based on the conformity of motif alleles to the genome^60^. For this analysis, we considered only lead SNPs found in our eQTL study. TF enrichment values were calculated as the ratio of the observed SNP hits over the expected ones for each TF. We considered enrichment for each TF then using an FDR cutoff of 0.05. TF enrichment was completed separately for each culture condition using the respective eQTL SNPs and ATAC-seq peaks.

## RESULTS

### Transcriptional profiling of human aortic SMCs

We performed RNA sequencing of aortic SMCs derived from 151 ethnically-diverse healthy heart transplant donors (118 males and 33 females) to identify their transcriptional profiles. After quality control filtering, data analyses were performed on 139 and 145 samples cultured in the absence (quiescent) or presence (proliferative) of FBS, respectively (**Figure 1**). The number of expressed genes ranged from 18,637 in quiescent to 18,116 in proliferative SMCs (**Table 1**). Principal component analysis identified two distinct clusters of samples corresponding to the cells cultured in the two conditions (**Supplementary Figure 1A**). Further, 2,773 genes were differentially expressed (*P*_adj_< 1×10^−3^), including canonical SMC markers (*VCAM1, SMTN, ICAM1, TAGLN, CNN1, ACTA2, SPP1*) in agreement with the differences observed in the quiescent and proliferative state of SMCs (**Supplementary Figure 1B, Supplementary Table 1**). Consistent with response to atherogenic stress, extracellular matrix organization was among the top biological pathways identified in GO enrichment analysis of upregulated genes in proliferative versus quiescent phenotypes. In contrast, pathways that are associated with DNA replication and proliferation were repressed in the quiescent phenotype (**Supplementary Figure 1C**).

**Figure 1:**
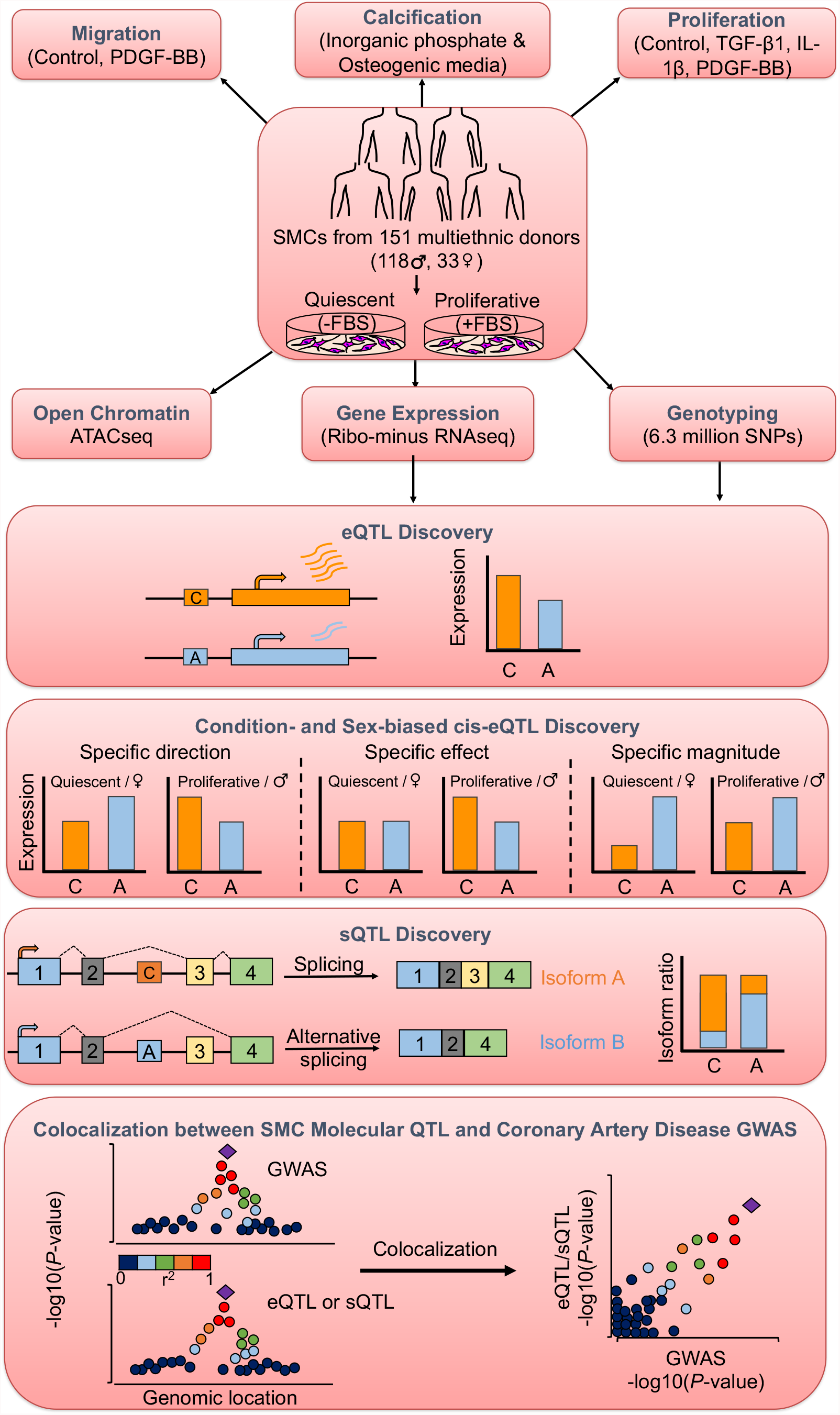
Study design and overview of analyses. Aortic smooth muscle cells (SMCs) from 151 multi-ethnic heart transplant donors were characterized for three atherosclerosis-relevant phenotypes: migration, calcification and proliferation^70^. To measure gene expression of SMCs, sequencing of ribosomal RNA-depleted total RNA isolated from SMCs cultured in the absence or presence of FBS to simulate quiescent or proliferative phenotypic state was performed. Associations of gene expression and splicing with the genotypes of ∼6.3 million imputed SNPs were calculated to discover *cis*-eQTLs as well as condition-specific and sex-biased eQTLs and sQTLs. Colocalization between molecular QTL and coronary artery disease GWAS associations was identified using four different methods.

To confirm that cultured SMCs reflect *in vivo* physiology, we projected their transcriptomes onto the 49 tissues profiled in GTEx v8^18^ (**Supplementary Figure 2**). We observed that SMCs formed a distinct cluster and closely neighbored fibroblasts, skeletal muscle, blood vessels and heart. These results suggest that cultured SMCs capture many aspects of in vivo physiology, thus supporting their utility in mapping candidate CAD genetic regulatory mechanisms.

### *cis*-Expression Quantitative Trait Loci in SMCs

We obtained genotype information for ∼6.3 million variants with at least 5% minor allele frequency in our population. Clustering of the donor genotypes with the 1000 Genomes reference population samples identified 6, 12, 64, and 69 of the individuals with East Asian, African, Admixed American, and European ancestry, respectively (**Supplementary Figure 3**).

To identify genetic loci associated with transcript abundance, we performed association mapping with the genotypes of ∼6.3 million variants and the expression levels of 18,116 and 18,637 genes in quiescent and proliferative SMCs, respectively using tensorQTL^41^. We identified 3,000 and 4,188 eGenes with *cis*-eQTL (< 1Mb) at FDR q-value < 0.05 in the quiescent and proliferative phenotypes, respectively (**Table 1**). We compared our results against the GTEx v8 eQTL dataset composed of 49 different human tissues^49^ using the QTlizer R package^48^. We found that 2,818 SMC *cis*-eQTLs (eSNP and eGene pair) were present in at least one GTEx tissue, whereas 3,660 were unique to our dataset (**Figure 2A**). Most of the shared SMC eQTLs were enriched in the tissues that are anatomically rich in SMCs (**Supplementary Figure 4)**. Conditioning on the lead SNPs identified 254 and 465 secondary and beyond eQTLs for quiescent and proliferative SMCs, respectively.

**Table 1:** Expression and Splicing Quantitative Trait Loci.

|  |  | Gene Expression |  | Splicing |  |
| --- | --- | --- | --- | --- | --- |
| Condition | Gene Type | No of genes tested | No of eGenes with at least one eQTL | No of genes tested | No of sGenes with at least one sQTL |
| **Quiescent** | protein coding | 13,341 | 2,267 | 11,123 | 2,774 |
|  | lncRNA | 3,183 | 477 | 959 | 244 |
|  | pseudogene | 1,484 | 215 | 291 | 129 |
|  | other | 629 | 41 | 1 | 0 |
|  | **Total** | **18,637** | **3,000** | **12,374** | **3,147** |
| **Proliferative** | protein coding | 13,182 | 3,367 | 11,013 | 3,180 |
|  | lncRNA | 2,986 | 546 | 996 | 273 |
|  | pseudogene | 1366 | 228 | 297 | 125 |
|  | other | 582 | 47 | 1 | 0 |
|  | **Total** | **18,116** | **4,188** | **12,307** | **3,578** |

**Figure 2:**
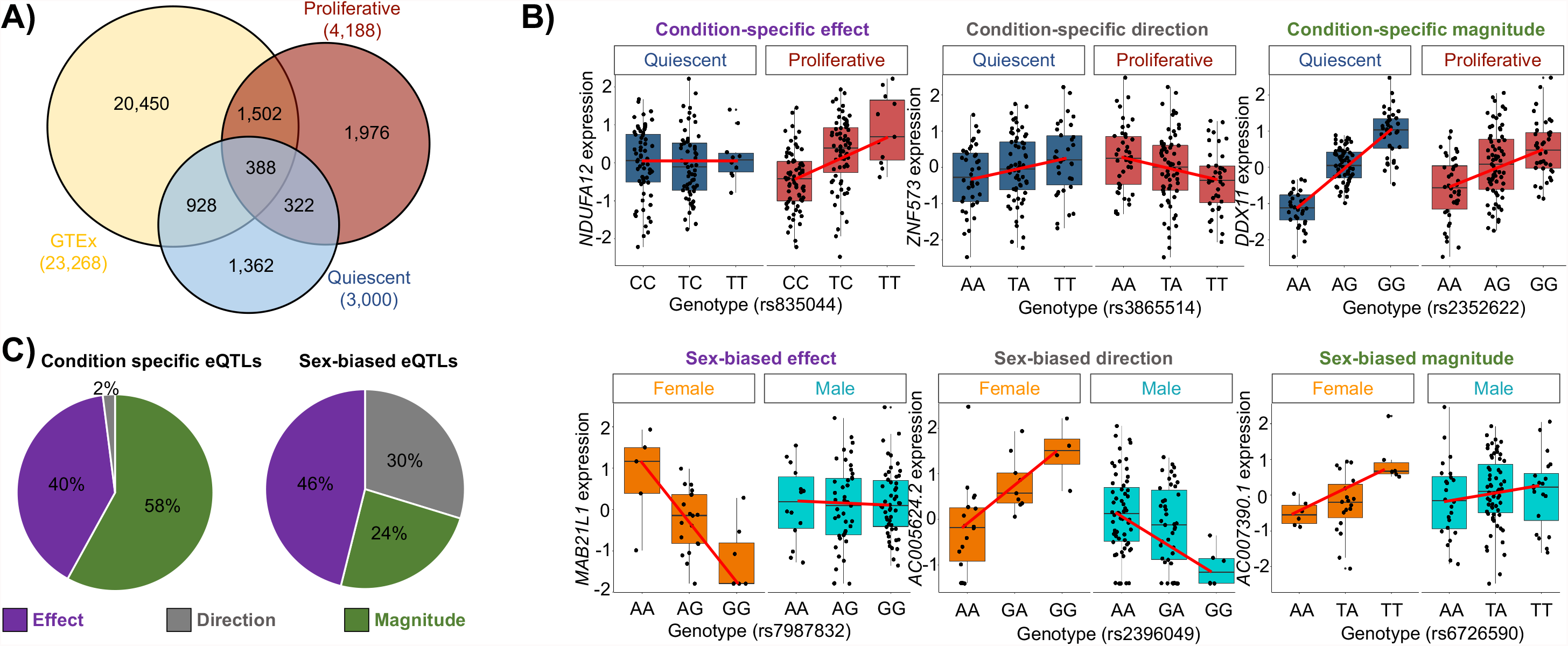
Identification of *cis*-eQTLs, condition-specific and sex-biased eQTLs in aortic smooth muscle cells. **A**) Venn diagram comparing eQTL discovered in quiescent (blue) and proliferative (red) conditions versus GTEx tissues (yellow) (FDR q-value<0.05). 3,660 of SMC *cis*-eQTLs (pair of SNP-gene) were absent or not significant in the GTEx dataset. 1,362 and 1,976 of these novel *cis*-eQTLs were unique to quiescent and proliferative cells, respectively. **B)** Condition-specific (top) and sex-biased (bottom) eQTLs were classified into three different categories: condition-specific or sex-biased effect, condition-specific or sex-biased direction, and condition-specific or sex-biased magnitude. **C**) Quantification of the three different condition-specific (left) and sex-biased (right) eQTL groups.

We also examined whether the SMC eQTLs present in the GTEx dataset^61^ are specific to SMCs using METASOFT^50^. We identified 29 SMC-specific *cis*-eQTLs under a stringent criteria of eQTL posterior probability > 0.9 for SMCs and < 0.1 for all the GTEx tissues (**Supplementary Table 2, Supplementary Figure 5**). To replicate/validate these results, we queried these 29 *cis*-eQTLs in the STARNET dataset of seven cardiometabolic tissues from ∼600 donors^19^. We identified only two of the SMC *cis*-eQTLs (rs367077-HLA-K and rs4795548-SH3GL1P2) to be present in the STARNET dataset at FDR < 0.05 (**Supplementary Table 5)**. 12 of the 29 SMC-specific *cis*-eQTLs were present in both quiescent and proliferative SMCs, 9 loci were *cis*-eQTLs only in quiescent and 8 loci were *cis*-eQTLs only in proliferative SMCs. Because many tissues in GTEx contain vascular wall cells, we examined if the 29 SMC eQTLs showed associations in monocytes/macrophages^62^ and aortic endothelial cells^20^. None of the 29 eQTLs were present in these cells suggesting that the regulatory impact of the variants in these loci are SMC specific.

15% and 39% of the eGenes were unique to quiescent and proliferative cells, respectively (**Figure 2A**). Therefore, we determined whether the eQTL effect sizes were statistically different between the two phenotypic states. We compared the regression slopes of an eQTL in quiescent (β_noFBS_) vs proliferative (β_FBS_) phenotypes using a Z-test^63^. We identified 1,248 eQTLs at FDR q-value < 0.05 with varying effects between the two phenotypic states (**Supplementary Table 3**). We classified the condition-specific eQTLs into three different categories (**Figure 2B**). 58% of them showed differences in magnitude of the effect size between the two phenotypes, 40% showed eQTL effect in only one phenotype, and 2% showed differences in the direction of effect between the two phenotypes (**Figure 2C**). Three examples of condition-specific eQTLs are shown in **Figure 2B**. This supports the idea that regulatory variation impacts SMC gene expression in specific contexts.

Since sex differences in the genetic regulation of gene expression have been observed in many tissues^64,65^, we separately identified sex-biased *cis*-eQTLs in quiescent and proliferative SMCs. We identified 457 and 454 sex-biased *cis*-eQTLs (eigenMT value <0.05) in quiescent and proliferative SMCs, respectively (**Supplementary Table 4**). 24% of them (combined conditions) showed differences in magnitude of effect size between the two sexes, 46% showed eQTL effect in only one sex, and 30% showed differences in the direction of effect between the two sexes (**Figure 2C**). Three examples of sex-biased eQTLs are shown in **Figure 2B**.

To characterize the potential function of the eQTL signals, we evaluated the overlap of the lead variant and LD proxies (R^2^>0.8; 1000G EUR) in accessible chromatin regions of SMCs as identified by ATAC-seq in 5 different donors in quiescent and proliferative phenotypes. Evaluation of the lead variants in these accessible chromatin regions overlapped ∼10% of the eQTL signals. After including LD proxies of these lead variants, 1,386 of 3,000 eQTL signals in quiescent SMCs and 1,945 of 4,188 eQTL signals in proliferative SMCs overlapped accessible chromatin regions, demonstrating the potential regulatory function of these loci. We next sought to identify transcription factor binding sites (TFBSs) overrepresented in ATACseq peaks and overlapping eQTL SNPs in the accessible chromatin regions (**Supplementary Figure 6)**. Motif enrichment analysis showed enrichment of putative binding sites for members of the SP2, SP1, ELK4, and GABPA TF families, some of which are known to play functional roles in SMC^66–68^.

### Colocalization between eQTLs and CAD GWAS signals

To predict the effector transcripts that are regulated by CAD GWAS loci, we performed colocalization analyses using four distinct approaches. Overall, we had genotypes for 169 of the 175 CAD loci in our dataset. First, to identify the eQTLs that were likely to be driven by the GWAS loci, we assessed if the GWAS and eQTL lead variants were in high linkage disequilibrium (LD) (R^2^ >0.8) in our population. Using this approach, we identified 16 and 22 eGenes (FDR q-value < 0.05) in the quiescent and proliferative phenotype that showed an overlap with CAD loci, respectively. We also performed colocalization analysis using three additional methods: SMR^51^, COLOC^53^, and eCAVIAR^13^. We used FDR q-value < 0.05 for SMR and colocalization posterior probability (CLPP) > 0.01 as cutoffs for eCAVIAR and PPH4 > 0.5 for COLOC. We identified 84 eGenes that showed statistically significant colocalization. 17 and 30 of them showed an overlap with at least two of the colocalization methods in quiescent and proliferative SMCs, respectively (**Figure 3, Supplementary Table 6, Supplementary Figure 7**). Some of the eGenes predicted by the four colocalization methods differed; therefore, we visualized the coincidence of the eQTL and GWAS lead SNPs by inspecting the regional colocalization plots using LocusCompare^54^. This coincidence was also supported by conditional analysis on each lead eSNP and CAD GWAS index SNP.

**Figure 3:**
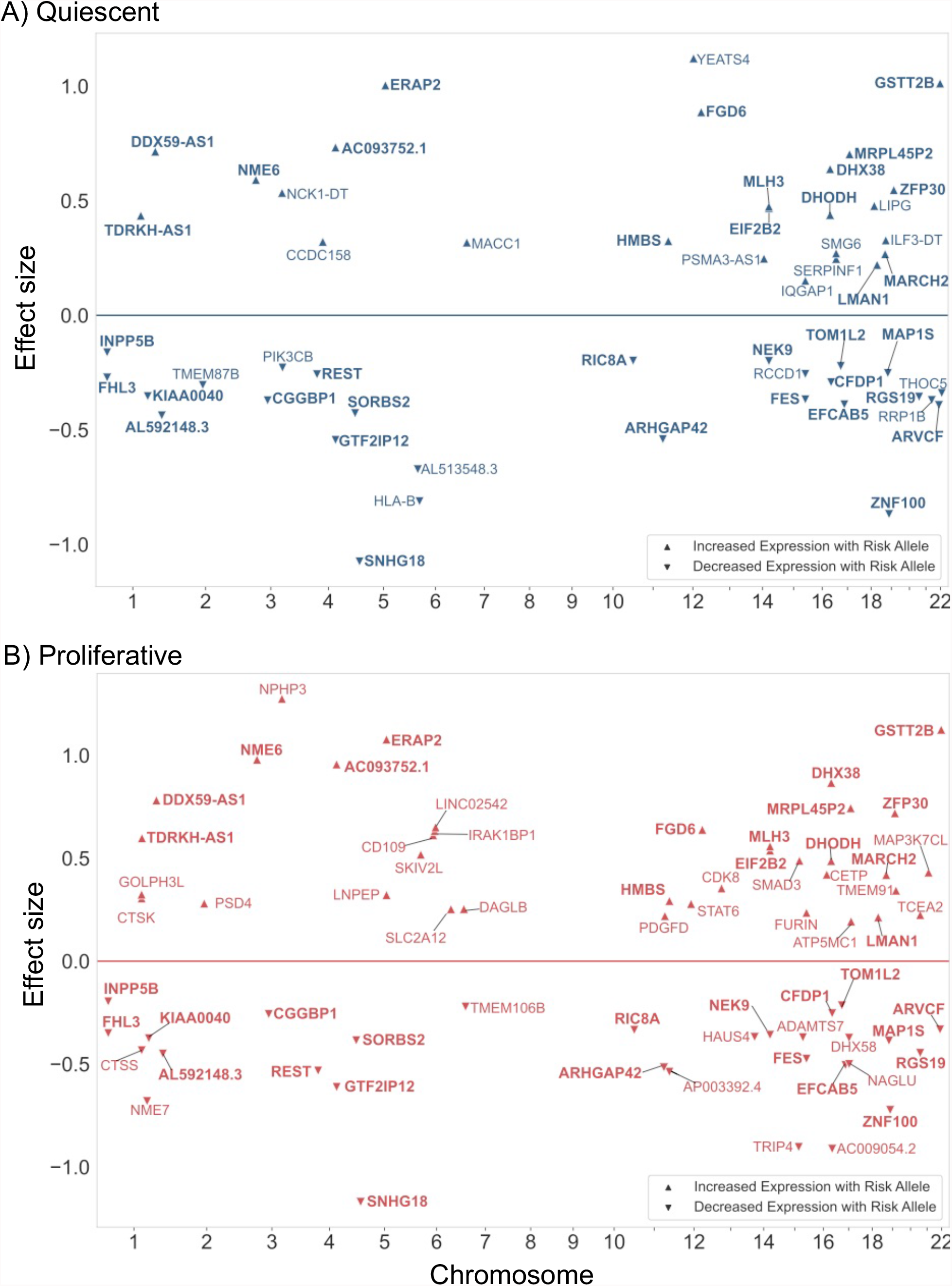
Summary of the SMC eQTL and CAD GWAS colocalization. Plots show the colocalization of CAD GWAS and eQTL signals using a combination of four different methods in **A)** quiescent and **B)** proliferative SMCs. The X-axis shows the chromosomal position of the colocalized SNP; Y-axis shows the effect size and direction of the eQTL with respect to the risk allele from the coronary artery GWAS^4^. Risk alleles that are associated with increased gene expression level are shown with up-triangle and risk alleles that are associated with decreased gene expression level are shown with down-triangle. Bolded gene symbols are common between the two phenotypic states.

Next, we assessed the LD between the sex-biased eQTL and CAD GWAS SNPs and identified two genes whose *cis*-eQTL colocalized with CAD loci: *AL160313*.*1* and *TERF2IP* (**Figure 4)**. In quiescent SMCs, *AL160313*.*1* had a *cis*-eQTL only in females; whereas, in proliferative SMCs, the *cis*-eQTL was in the opposite direction in males and females (**Figure 4A, top left**). Further, the CAD GWAS signal at this locus was stronger in males (P_rs11160538,male_=4×10^−4^; β_rs11160538,male_=-0.003) than in females (P_rs11160538,female_=1×10^−2^; β_rs11160538,female_=-0.001) in the UK Biobank cohort^69^ (**Figure 4B, bottom left**). In proliferative SMCs, the risk allele of the eSNP rs11160538, T, was associated with higher expression of *AL160313*.*1* in females and lower expression in males, suggesting a protective role for higher *AL160313*.*1* expression against atherosclerosis in males. The sex-biased eQTL for *TERF2IP* was present only in proliferative SMCs. The CAD risk allele of the eSNP rs12929673, T, was associated with lower expression of *TERF2IP* in females and higher expression in males (**Figure 4A, top right**). The CAD GWAS signal at this locus was stronger in males (P_rs12929673,male_=2×10^−6^; β_rs12929673,male_=0.28) than in females (P_rs12929673,female_=1×10^−2^; β_rs2929673,female_=0.15) in the UK Biobank cohort^69^ (**Figure 4B, bottom right**), suggesting a protective role for lower *TERF2IP* expression against atherosclerosis. The same locus was also associated with two SMC phenotypes relevant to atherosclerosis in a sex-stratified manner^70^. The CAD risk allele of the eSNP rs12929673, T, was associated with reduced proliferation response to IL-1β stimulation and lower calcification in females compared to males which showed the opposite effect (**Supplementary Figure 8**).

To identify SMC-specific genetic regulation of CAD risk, we asked if the SMC-specific eQTLs (**Supplementary Table 4**) colocalized with CAD GWAS loci. We found that *cis*-eQTL for *ALKBH8* colocalized with the 11q22.3 CAD locus (**Figure 5A**). This SMC-specific eQTL is also colocalized with systolic and diastolic blood pressure (**Figure 5B-C**), suggesting a role for this gene, which encodes a methyltransferase, in regulating blood pressure and atherosclerosis risk. The risk allele, G, of the eSNP rs7926602 is associated with lower expression of *ALKBH8* in quiescent and proliferative SMCs (**Figure 5D-E**). Re-analysis of gene expression data of atherosclerotic plaques obtained at carotid endarterectomy (GEO accession number GSE120521) showed that *ALKBH8* expression is reduced in unstable lesions compared to stable lesions (**Figure 5F**), implicating a protective role for higher SMC *ALKBH8* expression against hypertension and atherosclerosis.

**Figure 4:**
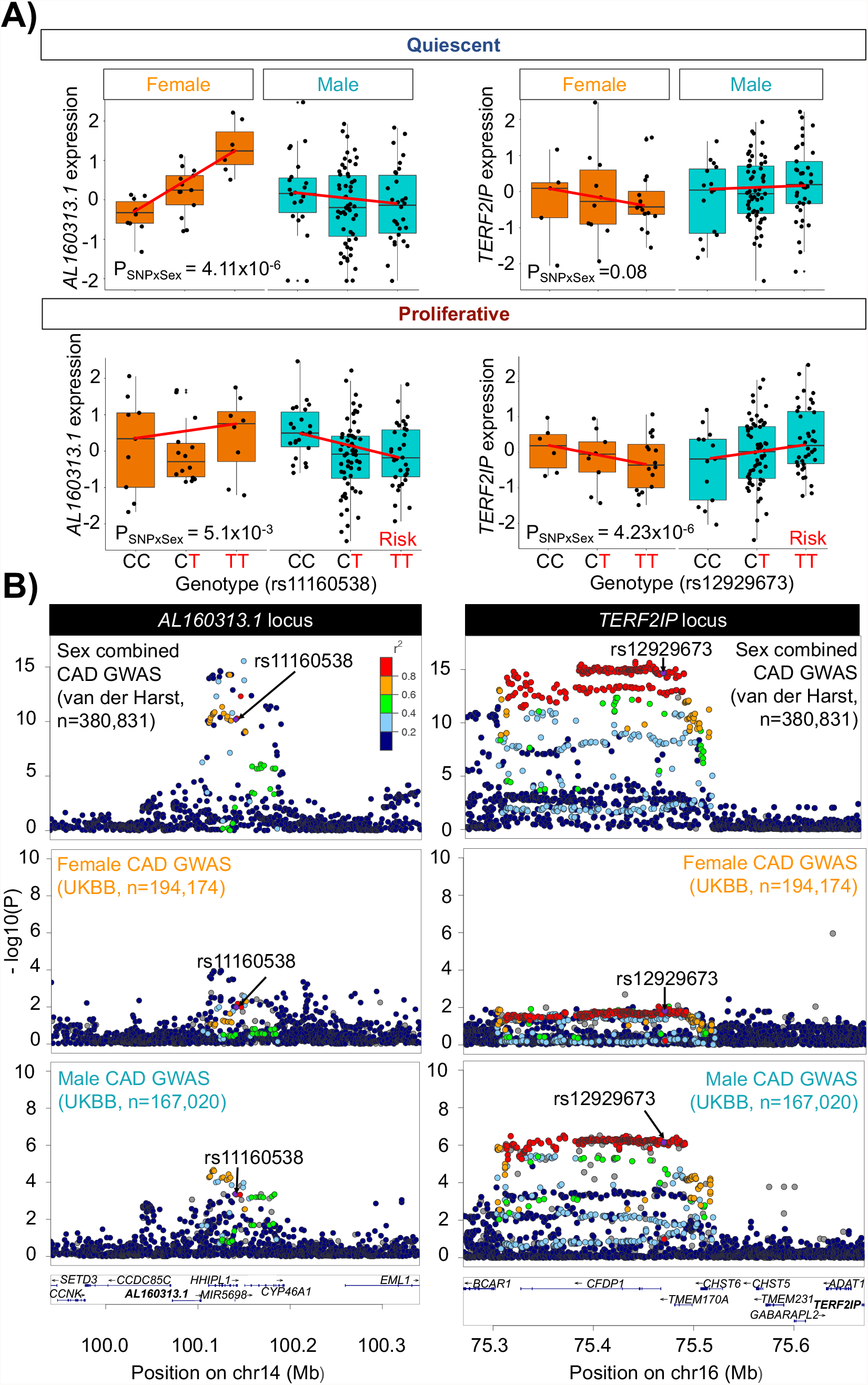
Colocalization between sex-biased-eQTLs and CAD GWAS. Colocalization based on linkage disequilibrium between sex-biased-eQTL SNPs and CAD GWAS identified two colocalized genes: *AL160313*.*1* and *TERF2IP*. **A)** Genotype-gene expression plots for *AL160313*.*1* (left) and *TERF2IP* (right) for the colocalized SNPs in quiescent and proliferative SMCs **B)** LocusZoom^45^ plots showing the association signal for sex-combined (Harst and Verweij^4^) and sex-stratified (UK Biobank^69^) GWAS for coronary artery disease at the *AL160313*.*1* or *TERF2IP* locus.

**Figure 5:**
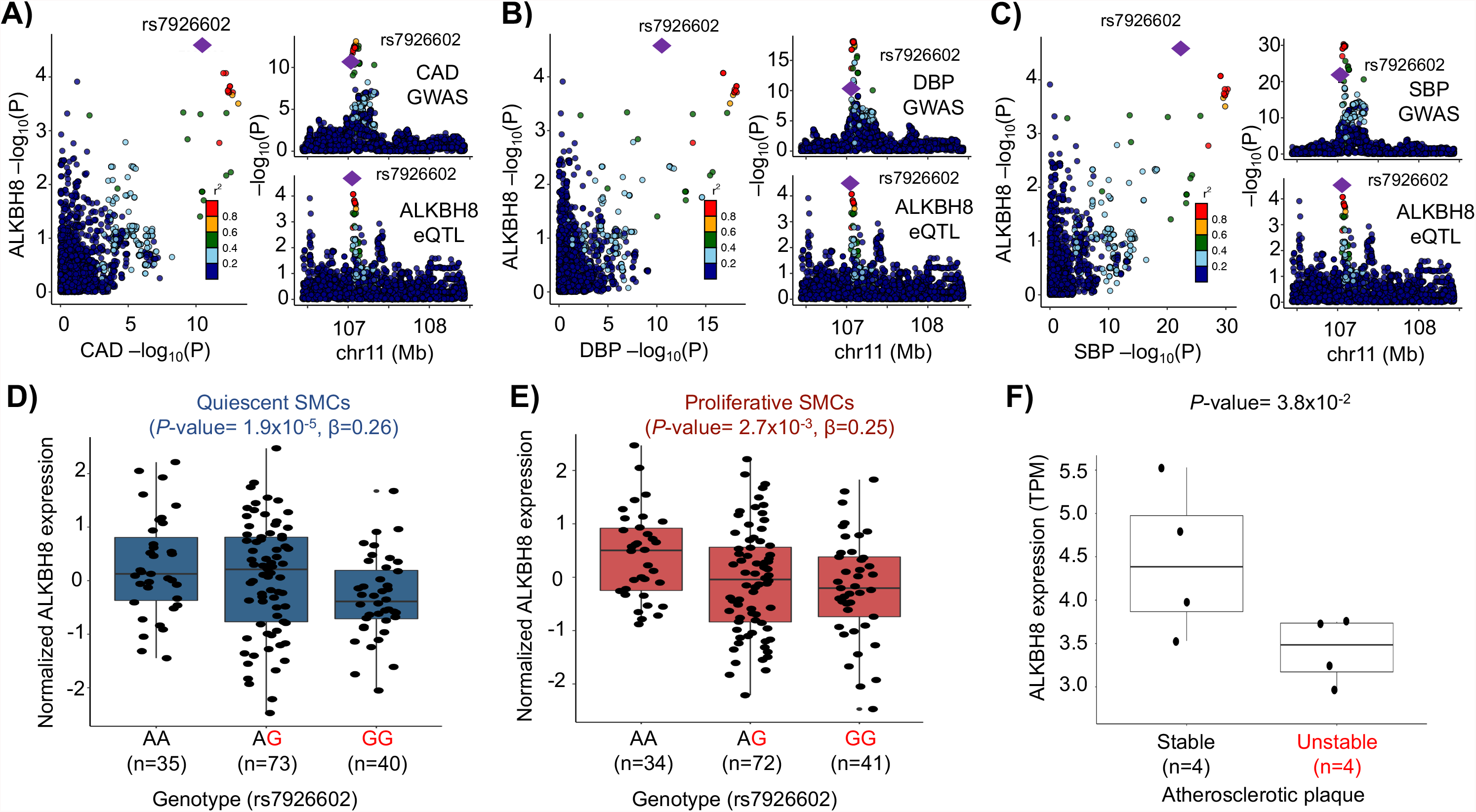
Colocalization between SMC-specific eQTL and vascular disease GWAS. *cis*-eQTL for *ALKBH8* expression colocalized with the 11q22.3 **A)** coronary artery disease (CAD), **B)** Diastolic (DBP) and **C)** systolic (SBP) blood pressure GWAS locus. The risk allele (G) of SNP rs7926602 is associated with lower *ALKBH8* expression in **D)** quiescent and **E)** proliferative SMCs. **F)** ALKBH8 expression in human unstable atherosclerotic plaques of carotid arteries^73^.

### Functional annotation of CAD GWAS-colocalized SMC eQTLs

The 84 colocalized eQTLs contain 3811 SNPs that are in high LD in our study population (R^2^>0.8). We overlapped these SNPs with accessible chromatin regions identified in ATACseq experiments performed in quiescent and proliferative SMCs from five donors. We determined that 128 SNPs in 30 eQTL loci in quiescent SMCs and 140 SxNPs in 37 eQTL loci in proliferative SMCs were located in accessible chromatin peaks (**Supplementary Table 7**). We predict these SNPs to have a regulatory impact on eQTL gene expression and potentially be causal for SMC gene expression and CAD risk.

We previously characterized the SMC donors for 12 atherosclerosis-relevant phenotypes^70^. First, we assessed the association of the eSNPs in the CAD loci with these phenotypes. Second, we assessed the correlation between the phenotypes and the colocalized eQTL gene expression. The risk allele of the eSNP rs7195958 at the *DHODH* locus had higher association with SMC proliferation compared to the non-risk allele. The risk allele was also associated with higher expression of *DHODH* in quiescent SMCs. As would be predicted by these association results, there was a significant positive correlation between *DHODH* expression and SMC proliferation, suggesting that the 16q22 CAD locus regulates SMC proliferation by perturbing *DHODH* expression (**Supplementary Figure 9A)**. Similarly, the risk allele of the eSNP rs12817989 at the *FGD6* locus had lower association with SMC proliferation compared to the non-risk allele. The risk allele was associated with higher expression of *FGD6* in proliferative SMCs. As would be predicted by these association results, there was a significant negative correlation between *FGD6* expression and SMC proliferation, suggesting that the 12q22 CAD locus regulates SMC proliferation by perturbing *FGD6* expression (**Supplementary Figure 9B)**.

Finally, human single cell RNAseq (scRNAseq) analysis from coronary atherosclerotic plaques confirmed the expression of most of the eQTL genes in SMCs (**Supplementary Figure 10**)^71^. We were able to assess the expression of 76 of the 84 eQTL genes in the coronary artery scRNASeq dataset and found that 50 were higher expressed in SMCs, pericytes, and fibroblasts compared to endothelial cells, monocytes, macrophages, and other immune cells. For example, we found that the expression of the long non-coding RNA SNHG18 was regulated by the 5p15 CAD locus in both the quiescent and proliferative SMCs. The risk allele was associated with decreased expression of SNHG18 (**Figure 6A**). Of the 16 potentially causal SNPs in high LD in this locus, six of them were located in accessible chromatin regions in both SMC phenotypic states (**Figure 6B, Supplementary Table 7**). Nine SNPs also displayed an allelic effect in a massively parallel reporter assay (MPRA) performed in SMCs exposed to cholesterol to induce phenotypic switching to resemble modulated SMCs found in atherosclerotic plaques^72^ (**Figure 6C**). SNPs rs1651285, rs1706987, and rs1398337 were in both accessible regions and showed allelic effects in MPRA, identifying them as the potential causal variants at this locus. The expression of *SNHG18* was highly enriched in SMC, fibroblast/fibromyocyte, and pericyte clusters in a human carotid artery scRNA-seq dataset^71^ (**Figure 6D**) and was downregulated in unstable atherosclerotic lesions compared to stable lesions from carotid arteries^73^ (**Figure 6E**). Further, SNHG18 expression was negatively correlated with PDGF-BB-induced proliferation (**Figure 6F**). When we silenced the SNHG18 expression in immortalized human coronary aortic SMCs (**Supplementary Figure 11**), we observed increased proliferation (**Figure 6G**). Collectively, these lines of evidence point to three variants in the 5p15 locus as potential causal SNPs regulating the expression of SNHG18 and SMC proliferation, thereby affecting the CAD risk in this locus.

**Figure 6:**
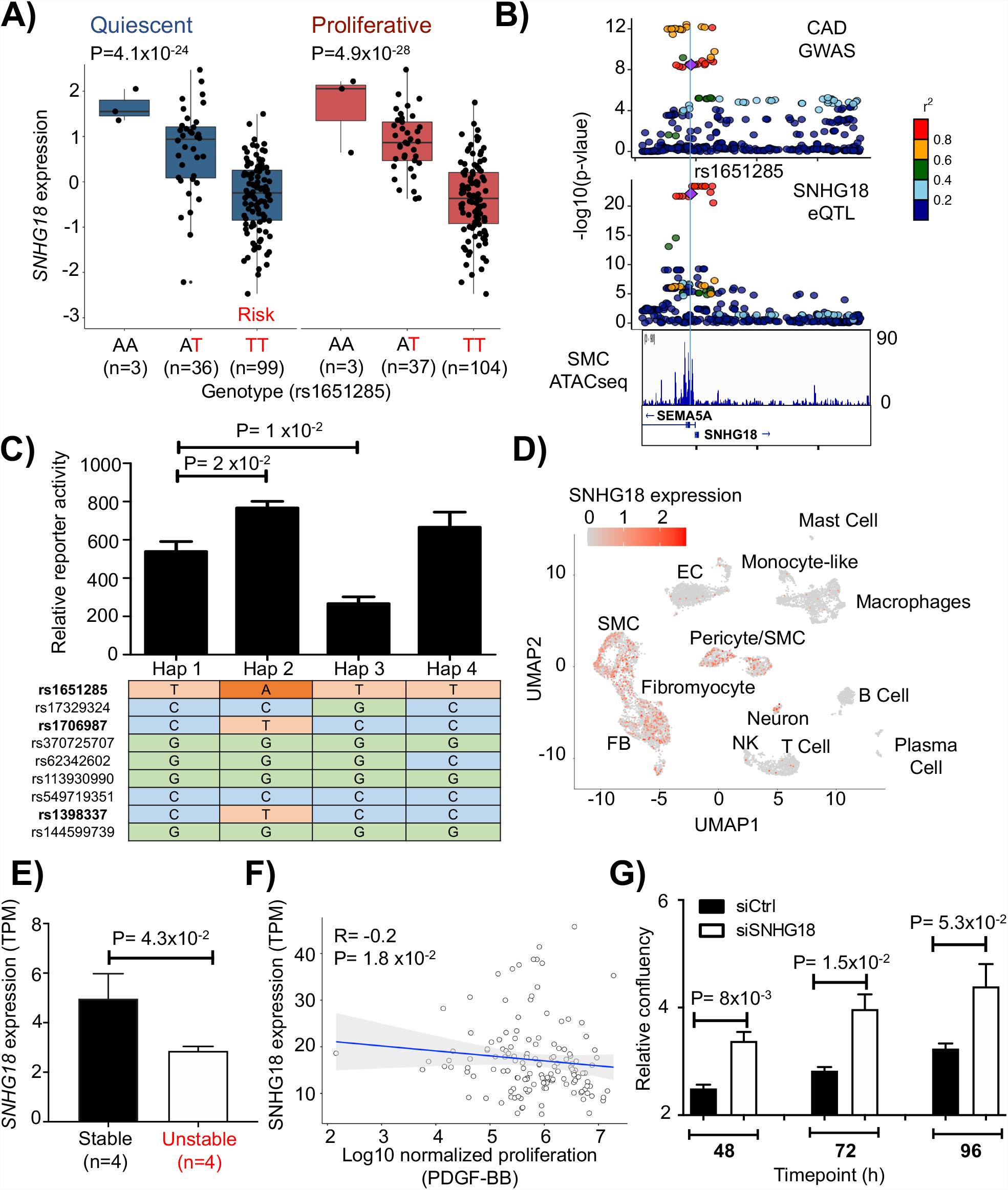
Colocalization between *SNHG18* eQTL and 5p15.31 CAD GWAS locus. The risk allele (T) of SNP rs1651285 is associated with lower *SNHG18* expression in **A)** quiescent and proliferative SMCs **B)** LocusZoom plots of the CAD GWAS and SMC *cis*-eQTL in the 5p15.31 locus. SNPs rs1651285, rs1706987, and rs1398337 are located in an accessible chromatin region identified by ATACseq in SMCs (lower panel). **C)** Bar plot summarizing the CAD haplotypes that demonstrated significant allele-specific enhancer activity in massively parallel reporter assays performed in cholesterol-loaded SMCs (n=3). SNPs located in an accessible chromatin region identified by ATACseq in SMCs are bolded. **D)** Uniform manifold approximation and projection plot of single-cell RNA-sequencing data from human coronary atherosclerotic plaques^71^. **E)** *SNHG18* expression in human stable and unstable atherosclerotic plaques of carotid arteries^73^. **F)** Negative correlation of *SNHG18* expression with PDGF-BB-induced proliferation in SMCs. **G)** Downregulation of SNHG18 in SMCs increased proliferation. Hap : haplotype.

### Splicing Quantitative Trait Loci in SMCs

Previous studies showed that RNA splicing explains a large proportion of heritable risk for complex diseases^9^. To identify genetic loci associated with mRNA splicing, we quantified RNA splicing with LeafCutter^43^ and performed association mapping with tensorQTL. We identified 3,147 and 3,578 sGenes with *cis*-sQTL (< 200 kb from splice sites) in the quiescent and proliferative phenotypes, respectively (FDR q-value <0.05) (**Table 1**). Similar to eQTL findings, the majority of the sGenes were shared (51%) between the two conditions. 19% and 29% of the sGenes were unique to quiescent and proliferative conditions, respectively (**Supplementary Figure 12A**). Conditioning on the lead SNPs identified 144 and 120 secondary and beyond sQTL genes for quiescent and proliferative conditions, respectively. We also determined the overlap of genes with *cis*-eQTL or -sQTL. In quiescent SMCs, only 20% of the 5,139 genes with a *cis*-eQTL or -sQTL were genetically regulated both at the mRNA splicing or expression levels. Similarly, in proliferative SMCs, only 24% of the 6,244 with a *cis*-eQTL or -sQTL were genetically regulated both at the mRNA splicing or expression levels (**Supplementary Figure 12B**). This suggests that genetic regulation of mRNA abundance and splicing are largely independent in SMCs, in agreement with studies in other tissues^9,74^.

### Colocalization between sQTLs and CAD GWAS signals

To identify genes whose alternative splicing is associated with genetic risk for CAD, we performed colocalization analyses of splicing QTLs and CAD loci using four distinct approaches similar to the eQTL analysis described above. The intronic excision levels, as measured by LeafCutter^43^, of 100 and 120 sGenes in the quiescent and proliferative phenotypes were significantly associated with CAD loci, respectively. Colocalization of *cis*-sQTLs with 44 and 60 genes with CAD loci was unique to quiescent or proliferative SMCs, respectively (**Figure 7A**; **Supplementary Table 8**). Significantly more CAD genes were colocalized with sQTLs (164) than eQTLs (84). We identified 11 genes whose expression and alternative splicing was associated with the same CAD loci (**Supplementary Table 9**). These results point to the significant role of the genetic regulation of mRNA splicing as a molecular mechanism for CAD genetic risk.

**Figure 7:**
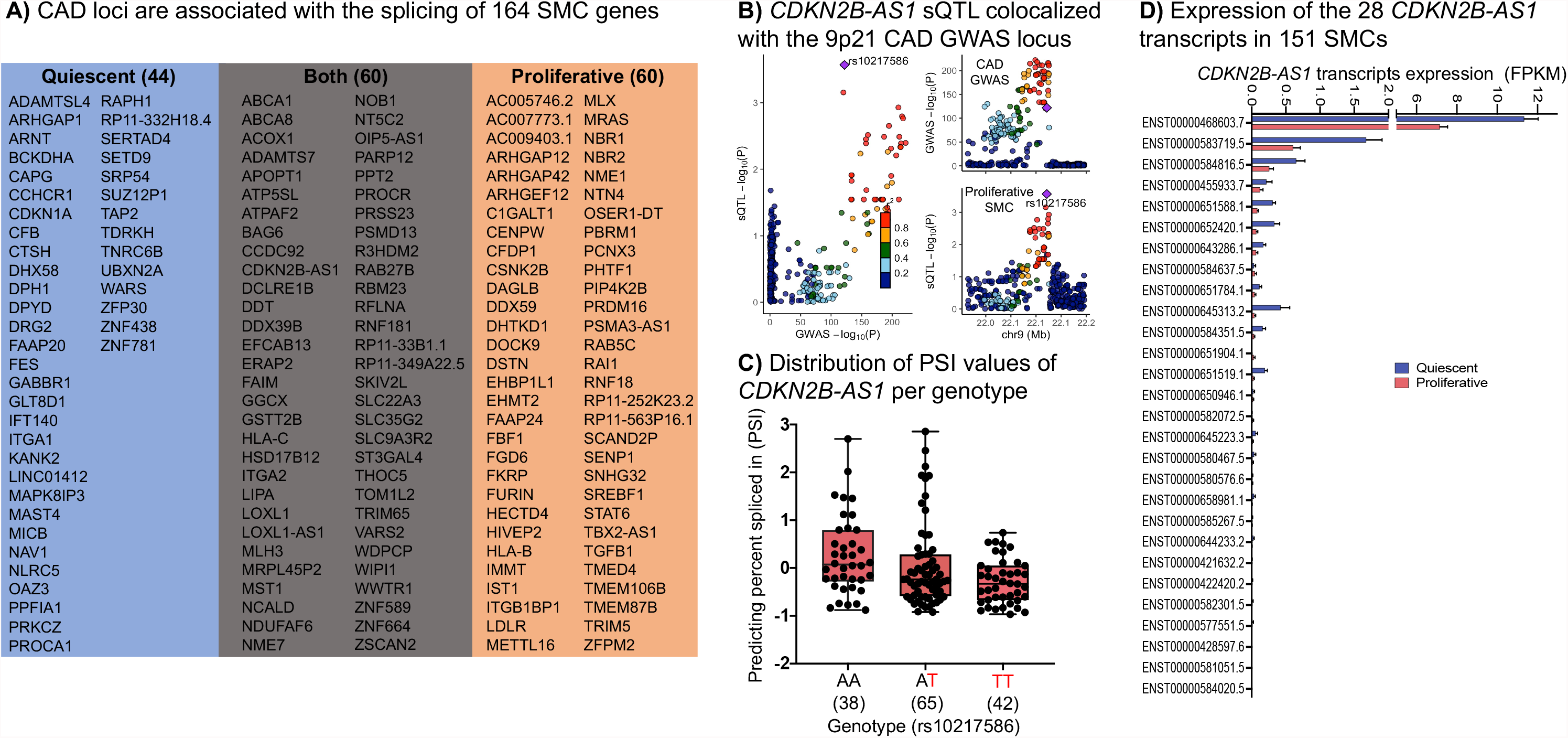
Summary of the SMC sQTL and CAD GWAS colocalization and the 9p21 locus. **A)** Shows the colocalization summary of CAD GWAS and sQTLs signals using four different methods in SMCs. **B)** *CDKN2B-AS1 cis*-sQTL signal colocalized with the 9p21 CAD GWAS locus. **C)** Distribution of the PSI values for CDKN2B-AS1 intron based on the rs10217586 genotype. **D)** Average expression of the 28 *CDKN2B-AS1* transcripts in 151 quiescent and proliferative SMCs. The standard error of the mean is shown.

We observed that the 9p21 locus, which has been the most significantly associated CAD locus in many populations, contains an sQTL for *CDKN2B-AS1*, also known as ANRIL (**Figure 7A-C**). We detected the expression of 25 of the 28 *CDKN2B-AS1* transcripts in SMCs (**Figure 7D**). Our results showed that the most significantly differentially excised intron at 9p21 (chr9:22064018-22096372) was found in *CDKN2B-AS1*. The frequency of this splicing event found in SMC proliferative phenotype (P = 2.7×10^−4^; β=-0.28) was colocalized with the genotype of the rs10217586 SNP in the CAD locus (P = 1.9×10^−122^; β=-0.16). Previous studies conducted in aortic endothelial cells^20,21^, monocytes^22^, whole blood^23^, and coronary artery SMCs^24^ did not identify an eQTL or sQTL for *CDKN2B-AS1* in this locus, suggesting that the genetic variants in the 9p21 locus act in aortic SMCs through splicing.

## Discussion

GWAS have successfully identified more than 200 loci associated with CAD risk; however, the genes and mechanisms responsible for many of these loci remain unknown. Majority of the variants are in non-coding regions making the task of identifying causal variants and genes difficult. Systems genetics aims to address this challenge by associating genetic variants with molecular phenotypes to comprehensively uncover the relationship between genotype and phenotype. Colocalized molecular QTL signals enable identification of reasonable candidate genes in disease-relevant cells and tissues. Therefore, we conducted, as far as we know, the largest transcriptome and whole-genome analyses using human aortic SMCs derived from a multiethnic population. We cultured these critical vascular cell types to CAD in two different media formulations to recapitulate disease and healthy conditions. PCA analysis of the transcriptome confirmed the distinctiveness of the two conditions and the comparison with publicly available datasets in GTEx revealed regulatory patterns specific to human SMCs.

Intersection of our SMC eQTL data with the GTEx dataset indicated that more than half of SMC eQTLs were not evident in GTEx tissues, indicating genetic regulation of gene expression unique to SMCs. Another study also found about half of the eQTLs from aortic endothelial cells of up to 157 donors were absent in the GTEx dataset^21^. This is possibly because most GTEx eQTLs have been performed in heterogeneous tissue samples containing various cell types and the genetic effects that are functioning only in rare cell types within a sampled tissue may not be detected. Indeed, most of the SMC eQTLs shared with GTEx samples were in the tissues that are rich in SMCs. Differences in RNA sequencing methods may have also contributed to the differences between GTEx and our study. Cell-type-specific eQTL analysis in disease-relevant tissues will lead to the identification of novel and more precise disease associations that can help elucidate the molecular mechanisms by which the genetic variants affect the disease.

Overlapping transcription factor binding sites with eQTL SNPs identified enrichment of putative binding sites for members of the SP2, SP1, ELK4, and GABPA TF families. While SP2 has an unknown role in SMCs, we predicted that the eQTL SNPs would impact SP2 binding to DNA in both the quiescent and proliferative phenotypes, suggesting an important role for this transcription factor in the regulation of gene expression by genetic variants in SMCs. We also predicted the transcription factors KLF5, E2F1, and CTCF to be important for disease development as their binding sites are enriched only in the proliferative SMCs. E2F1 is not only known to regulate cell proliferation^101^ but it is also upregulated in proliferative SMCs compared to quiescent SMCs suggesting it may be contributing to the phenotypic transition of SMCs from a quiescent to a proliferative state.

Several colocalization approaches have been developed in recent years^75^. They are sensitive to the parameters, such as thresholds applied to the prior probabilities, and differences in haplotype structures of the populations from which GWAS and molecular QTL data are derived. When the lead variants for the GWAS and eQTL studies are the same or in high LD in both populations, colocalization is straightforward^76^. Since our population had multi-ethnic ancestry and the CAD GWAS participants were of European ancestry, we used four different colocalization methods that may help to account for the differences in the LD structure. While multiple eQTLs and sQTLs had evidence for colocalization with CAD loci with two or more approaches, only seven genes (*AL513548*.*3, EIF2B2, FES, FURIN, MAP3K7CL, SMAD3, REST*) had evidence from all four approaches. Further development of colocalization methods are needed to increase the confidence in molecular QTL studies for identifying candidate genes for GWAS loci.

A previous study predicted the colocalization of five genes with CAD loci using eQTL data from coronary artery SMCs of 52 donors cultured only under proliferative conditions^24^. Our study of 151 donors whose SMCs were cultured in two conditions significantly expands these previous findings. Only two of the five predicted causal genes in the coronary SMC study, *FES* and *SMAD3*, were replicated in our study. One of the five genes, *TCF21*, which encodes a transcription factor that inhibits SMC differentiation^71,77^, is expressed in coronary but not aortic SMCs; therefore, we were not able to test its association with genetic variants. While the other two genes, *SIPA1* and *PDGFRA*, are expressed in aortic SMCs, we did not detect their colocalization with CAD loci. The other discrepancies between the two studies may be related to differences in methods, such as the *P*-value thresholds for declaring a *cis*-eQTL or its colocalization significant. By performing our studies in a larger number of donors with deeper RNA sequencing, we significantly increased the number of the predicted causal CAD genes playing a role in SMCs. The differences between aortic and coronary artery SMC eQTLs may point to differences in transcriptional regulation between the two vascular beds. Larger numbers of donors or meta-analysis of the SMC eQTL studies should lead to the identification of more causal genes associated with CAD.

Nearby genes to the index SNP in a locus are usually used as signposts for annotation. Previous CAD GWAS identified 366 nearby genes as potentially causal^4^. We found that only 26 and 33 of these nearby genes matched eQTL and sQTL, respectively. On the other hand, 173 genes were distinct from the initial locus annotations. 58 were derived from eQTL and 127 were derived from sQTL colocalizations. For example, rs11810571 in the 1q21.3 locus is located near *TDRKH* and *RP11-98D18*.*9*. However, our results showed significant colocalizations with the expression level of *GOLPH3L, CTSK* and *CTSS* located ∼ 1 MB from the variant. For 8 loci, we identified multiple genes associated with the risk variants. For example, 15q21 risk locus was associated with *FES* and *FURIN* genes in proliferative state, while in quiescent SMCs it was associated with the expression of *FES, RCCD1* and *IQGAP1*. Finally, previous GWAS identified missense mutations in 20 genes^4^. For two of the genes, *LIPA* and *TRIM5*, we also observed an eQTL effect. For three of the genes, *ADAMTS7, DAGLB, DHX58*, we also observed both eQTL and sQTL effects. These examples demonstrate the complexity of the molecular mechanisms by which CAD loci affect disease risk.

Long non-coding RNAs (lncRNAs) are typically >200 nucleotides in length and do not contain a functional open reading frame. They can be encoded within protein coding genes or can be encoded in the intergenic regions from the sense or antisense DNA. They are expressed at much lower levels relative to their protein coding counterparts^78^. By performing library preparation with ribosomal RNA depletion, as opposed to polyA selection, and deep sequencing, we were able to assess the expression of ∼3,000 lncRNAs. Recent studies have shown that lncRNA plays an essential role in SMC biology and CAD^79,80^. We identified that CAD loci were associated with the expression of 12 lncRNAs and with the splicing of 15 lncRNAs. One of the colocalized lncRNA was small nucleolar RNA host gene 18 (*SNHG18*) that was regulated by the variants in the 5p15 CAD locus in both the quiescent and proliferative SMCs. While the role of SNHG18 in SMC biology and CAD have not been studied, it was observed to be upregulated in glioma and regulate the progression of epithelial-mesenchymal transition and cytoskeleton remodeling of glioma cells^81^.

Identifying tissue and cell-specific mechanisms of GWAS loci has been challenging with few notable exceptions ^82–84^. We found that SMC-specific eQTL for *ALKBH8* colocalized with the 11q22.3 CAD locus, with the risk allele leading to lower *ALKBH8* expression. The CAD risk allele is also associated with higher blood pressure^69^, suggesting a role for this gene, which encodes a tRNA methyltransferase, in regulating the vascular tone. Embryonic fibroblasts isolated from *Alkbh8*-deficient mice were shown to have increased levels of intracellular reactive oxygen species (ROS), lipid peroxidation products and a transcript expression signature indicative of oxidative stress compared to fibroblasts isolated from wild-type littermates^85^. *Alkbh8* deficient mice were characterized by abnormal sinus arrhythmia and decreased heart rate variability^86^. ALKBH8 agonists have been proposed for treating myocardial infarction injury due to its function on the modulation of autophagy and oxidative stress^87^.

Significant differences between sexes in the underlying pathology of atherosclerosis and its gene regulation have been described by us and others^17,88^. We had 118 male and 33 female donors in our population. The sex ratio of the donors is similar to the reported cases over a 20-year period in heart transplant donor registries, where 31.3% of the transplanted hearts are from deceased women^89^. Despite the imbalance in the numbers of males and females, which may have effected the statistical power of the interaction test, we were able to identify ∼1,000 sex-biased eQTLs. Colocalization analysis between sex-biased eQTL and GWAS summary statistics yielded loci that were either not detected or had a weak signal in sex-combined GWAS. This observation is consistent with previous findings^64^. Therefore, combining sex-stratified eQTLs with summary statistics from sex-stratified GWAS is necessary to understand the impact of sex on CAD driven by SMC dysfunction. One example of sex-biased eQTLs that colocalized with CAD GWAS signal was the *TERF2IP gene*. Lower *TERF2IP* leads to telomere elongation^90^, which is associated with decreased CAD risk^91^. Risk allele at this locus was associated with higher *TERF2IP* expression in males and lower expression in females. The same locus had an association with CAD risk only in males, suggesting that the lower *TERF2IP* expression in females may be playing a protective role against atherosclerosis.

Identifying genes whose expression is influenced by colocalizing *cis*-eQTL is just the first step in dissecting SMC and CAD GWAS loci. Discovering the functions of the predicted causal genes in SMC biology and CAD risk is also needed. We had previously shown that CAD loci are associated with atherosclerosis-relevant cellular phenotypes in the same donors^70^. We combined the two datasets to predict that *DHODH* and *FGD6* are candidate causal genes which regulate SMC proliferation. *DHODH* encodes dihydroorotate dehydrogenase, which catalyzes the fourth enzymatic step in *de novo* pyrimidine biosynthesis. Its role in SMCs is not known but pyrimidine nucleotides are involved in the energetics of smooth muscle contracture^92^. A missense variant in *FGD6* has been shown to increase risk of polypoidal choroidal vasculopathy, which primarily affects the vascular layer of blood vessels in the choroid^93^.

The 175 CAD loci contain >6,000 SNPs and identifying which of these alter transcriptional activity in SMCs is a necessary step to dissect the molecular mechanism of the loci. We overlaid the genomic locations of eQTL SNPs with accessible chromatin regions to predict that 194 SNPs may alter gene expression, thereby significantly reducing the number of predicted causal variants in CAD loci. Our studies show that integrating across multiple scales, from genotype to cellular phenotypes, allows us to focus on a few plausible hypotheses to test in subsequent *in vitro* and *in vivo* studies to identify the molecular and cellular genetic mechanisms of CAD loci.

The landscape of CAD-relevant RNA splicing events are mostly unknown. We observed that significantly more CAD loci were associated with splicing than expression, suggesting that the majority of the genetic risk for CAD acts through regulating transcript splicing in SMCs rather than transcript abundance. This observation is in agreement with a previous study that showed that sQTLs are more likely to be enriched for Alzheimer’s disease GWAS SNPs than eQTLs^74^. Further, we observed that sQTLs that colocalized with CAD loci were associated with a distinct group of genes than eQTLs, indicating that our results can explain additional factors of the genetic architecture of CAD.

Of significant note, we identified the colocalization of *CDN2B-AS1* (ANRIL) sQTL with the 9p21 locus. This locus has been under intense scrutiny because it is the most significantly associated CAD locus and has been replicated in many populations with diverse ancestries^94^. Since there is no association with traditional risk factors such as dyslipidemia, diabetes mellitus, age, and sex, previous studies focused on identifying the effects at the vessel wall. eQTL studies in endothelial cells did not identify an impact of the genetic variants in this locus on gene expression^21^; however, they have been shown to regulate adhesion, contraction, and proliferation in SMCs derived from induced pluripotent stem cells^95^. Aortic SMCs isolated from mice with a knock-out of the homologous region showed excessive proliferation and diminished senescence^96^. When these mice were bred to an atheroprone background, they developed larger atherosclerotic plaques with no changes in blood pressure, lipid levels, body weight, or fasting glucose^97^. Primary SMCs were prone to dedifferentiation and had accelerated calcification, reflective of the susceptibility mechanisms of the humans carrying the risk allele. This region contains five tightly clustered genes, which partly overlap. *CDKN2B-AS1*, also known as *ANRIL*, overlaps in antisense the full length of the *CDKN2B* gene body, while sharing a bidirectional promoter with *CDKN2A*. There are 28 linear and multiple circular isoforms of *ANRIL*. We detected the expression of 25 of the 28 linear isoforms in SMCs. Previous studies showed associations of linear *ANRIL* isoforms, as well as *CDKN2A* and *CDKN2B* with the variants in the 9p21 locus in whole blood, peripheral blood monocytes, peripheral blood T lymphocytes, lymphoblastoid cells lines, vascular tissues such as carotid atherosclerotic plaque samples, aorta, mammary artery, as well as subcutaneous or omental adipose tissue^94,98^. Our study shows these variants affect linear ANRIL splicing in SMCs; however, the associations of these variants with circular forms of ANRIL remain to be determined. The mechanism by which *ANRIL* isoforms affect SMC functions such as proliferation, migration, and calcification also needs to be explored.

Collectively, our results predicted candidate causal genes playing a role in SMCs that modulate the genetic risk for CAD. Some of the loci act differentially in quiescent and proliferative SMC phenotypes emulating different stages of atherosclerosis. They also have distinct effects in males and females and some are SMC-specific. Taken together, our results provide evidence for the complexity of the molecular mechanisms of CAD loci. We expect that our findings will provide a rich catalog of molecular QTLs to the cardiovascular community and candidates for future preclinical development.

## Data Availability

RNAseq data is available at GEO with the accession number GSE193817. eQTL/sQTL results can be accessed at https://virginia.box.com/s/t5e1tzlaqsf85z13o4ie2f9t1i0zfypd and https://virginia.box.com/s/o81cxrj5xne3xem4au785mupikduuwbu

### Web Resources

DESeq2, https://bioconductor.org/packages/release/bioc/html/DESeq2.html

FastQC, v0.11.5, https://www.bioinformatics.babraham.ac.uk/projects/fastqc/

King, 2.2.4, http://people.virginia.edu/~wc9c/KING/kingpopulation.html

LeafCutter, v0.2.9, https://github.com/davidaknowles/leafcutter

Matrix eQTL, v2.3, https://cran.r-project.org/web/packages/MatrixEQTL/index.html

METASOFT, http://genetics.cs.ucla.edu/meta_jemdoc/index.html

NGSCheckMate, v.1.0.0, https://github.com/parklab/NGSCheckMate

PEER, v1.3, https://www.sanger.ac.uk/science/tools/peer

PLINK, v1.90b6.16, https://www.cog-genomics.org/plink2/

qvalue Package, 2.14.1, https://www.bioconductor.org/packages/devel/bioc/html/qvalue.html

RNA-SeQC, v2.3.4, https://software.broadinstitute.org/cancer/cga/rna-seqc

Samtools, v1.10, https://samtools.github.io

STAR, v2.5.3a, https://github.com/alexdobin/STAR

sva, https://bioconductor.org/packages/release/bioc/html/sva.html

tensorQTL, v.1.0.2, https://github.com/broadinstitute/tensorQTl

Trim Galore,0.6.5, https://www.bioinformatics.babraham.ac.uk/projects/trim_galore/VerifyBamID, v1.1.3, https://genome.sph.umich.edu/wiki/VerifyBamID

### Data repositories

eQTL/sQTL results can be accessed at https://virginia.box.com/s/t5e1tzlaqsf85z13o4ie2f9t1i0zfypd and https://virginia.box.com/s/o81cxrj5xne3xem4au785mupikduuwbu

GENCODE v32, https://www.gencodegenes.org/human/

GTEx v8, https://www.gtexportal.org/home/datasets

1000 Genomes Phase 3, https://www.internationalgenome.org/category/phase-3/

