## Supplementary Figures for "Genetic regulation of human aortic smooth muscle cell gene expression and splicing predict causal coronary artery disease genes"

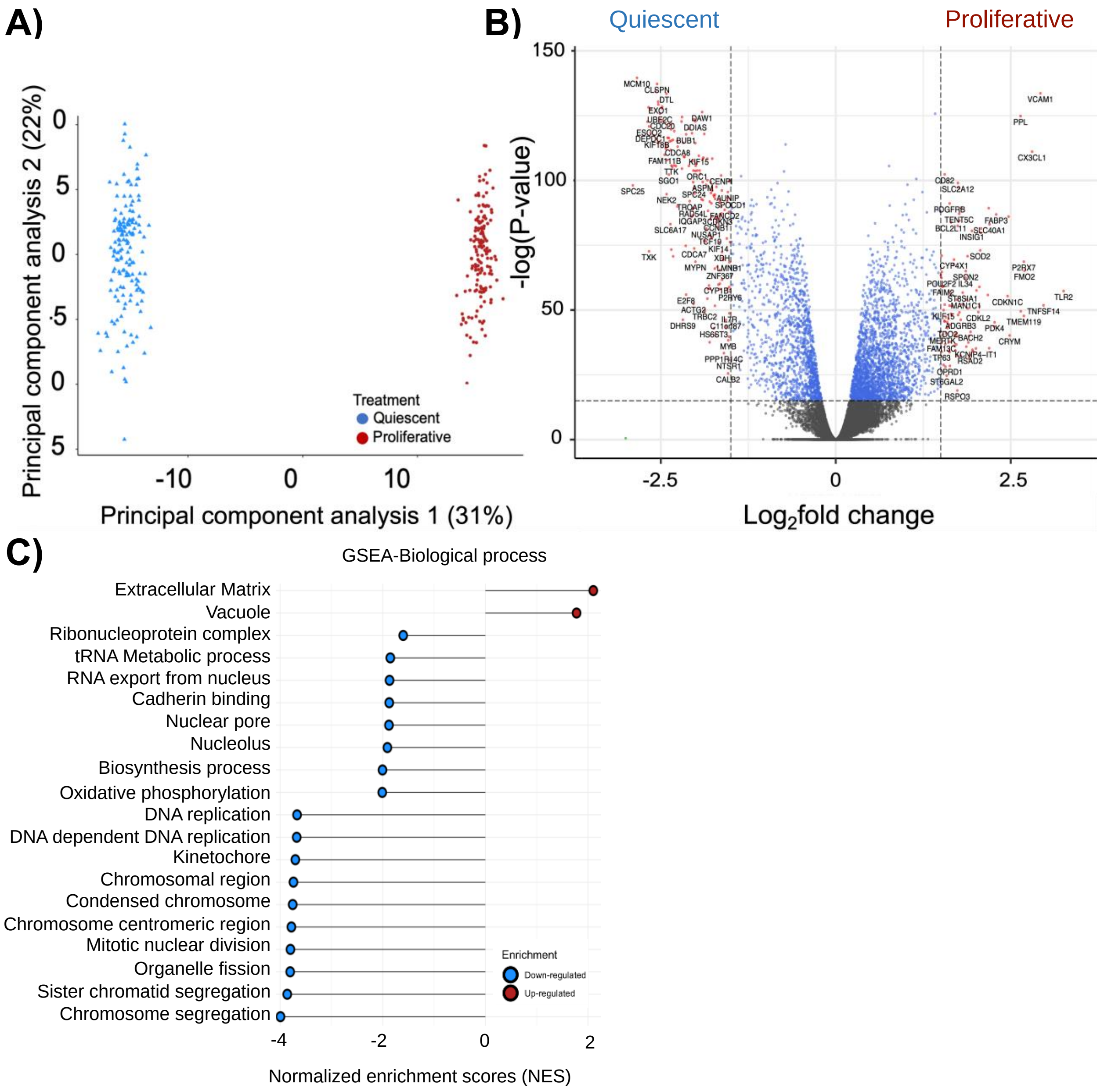


**Supplementary Figure 1: Transcriptional profiling of human aortic smooth muscle cells. A)** Principal component analysis of gene expression in quiescent and proliferative conditions. **B)** Volcano plot of the expression profiles of genes down and upregulated in SMCs cultured in quiescent and proliferative conditions. The red points represent the differentially expressed genes, gray points represent genes with no difference in their expression. The vertical dashed lines correspond to a 1.5-fold change in expression (up or down), and the horizontal dashed line represents the adjusted P-value ($P_{adj}$<0.05). **C)** Gene Ontology (GO) pathway analysis of the 2,773 differentially expressed genes, up-regulated (red) and down-regulated (blue) in quiescent and proliferative conditions.

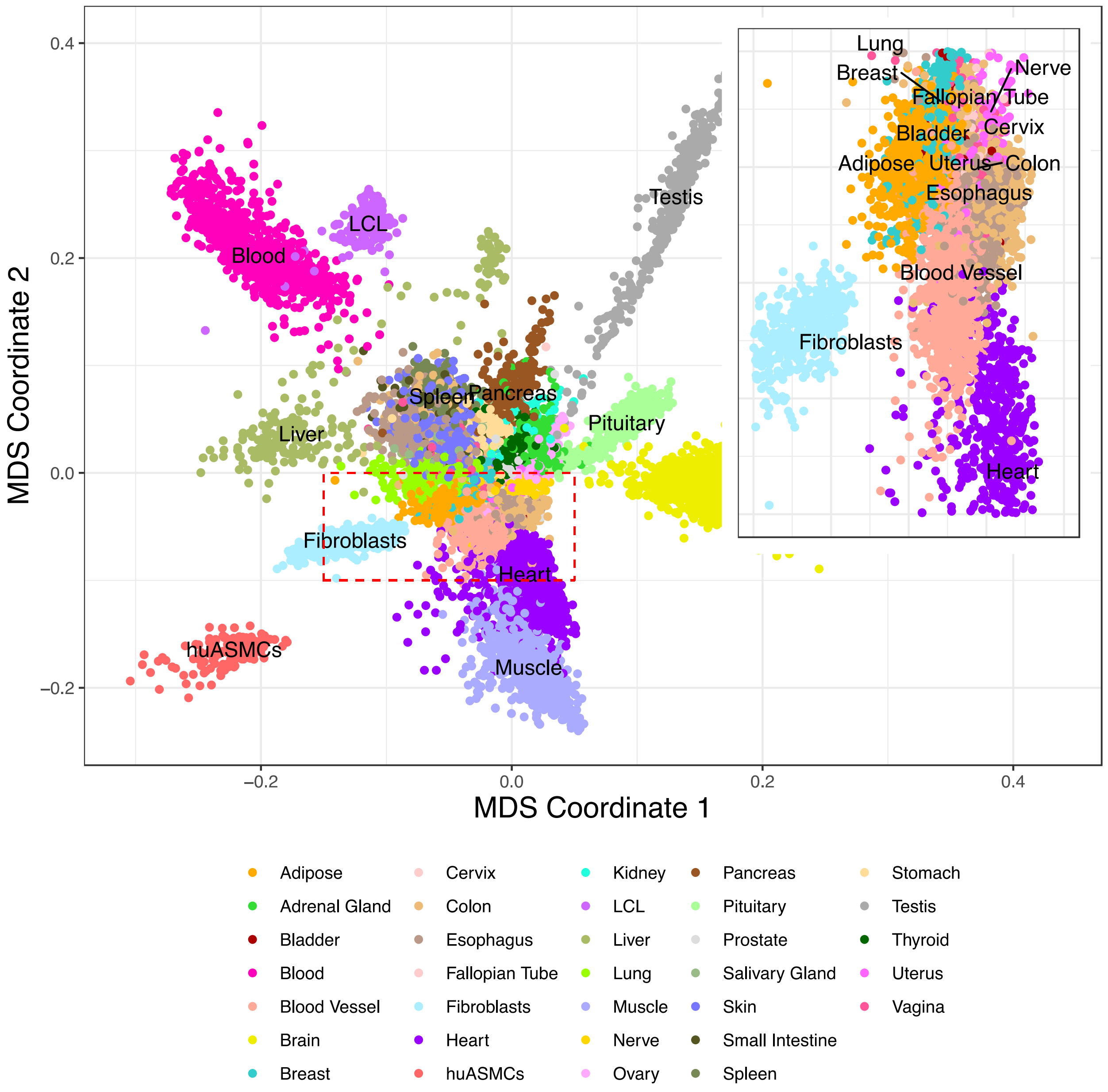


**Supplementary Figure 2: Comparison of aortic smooth muscle cell transcriptome with the transcriptomes of tissues and cells profiled in the Genotype-Tissue Expression (GTEx) project**. The multidimensional scaling plot of gene expression of SMCs and GTEx tissues and cell types shows a distinct cluster, which neighbors fibroblasts, skeletal muscle, blood vessels and heart (inset).

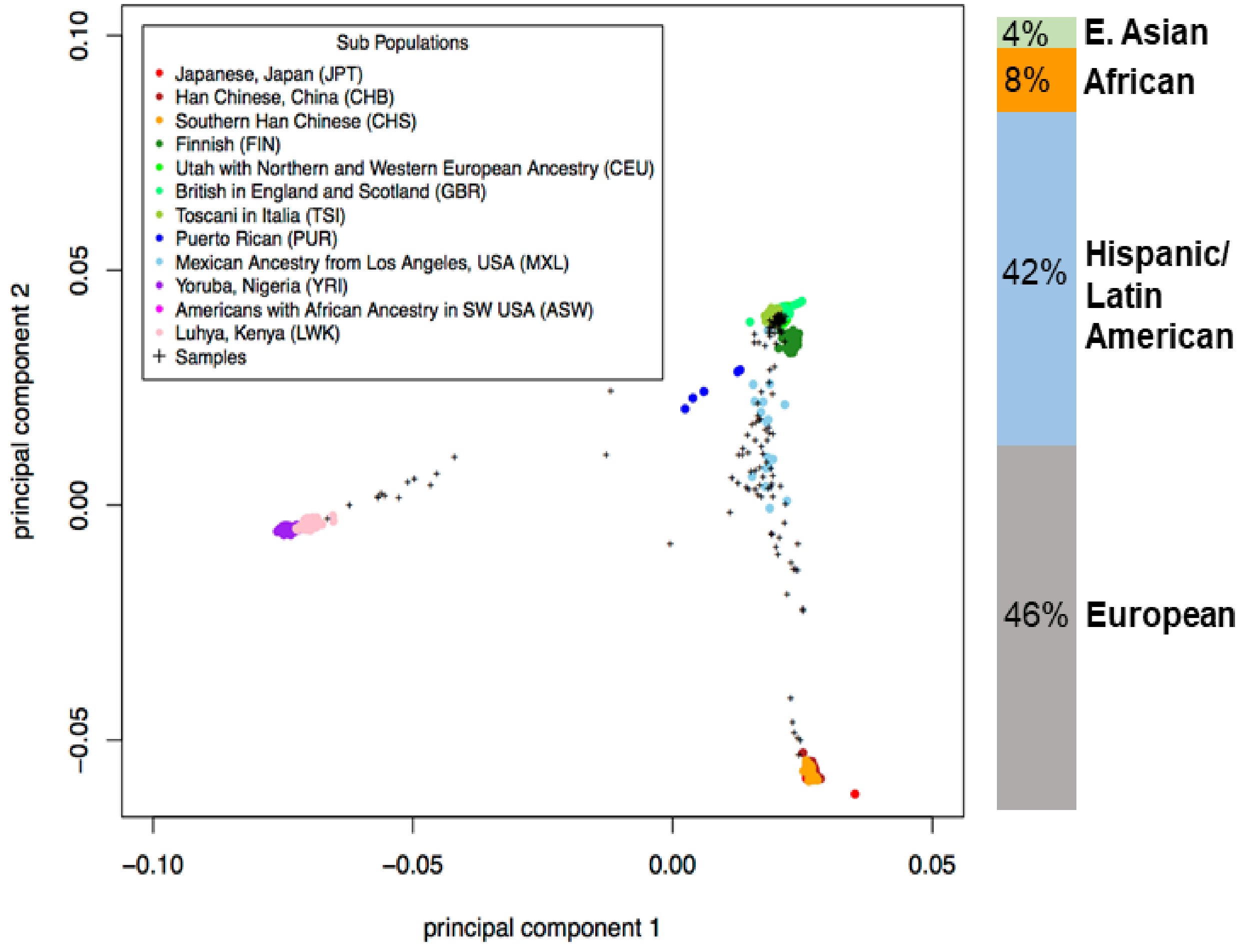


**Supplementary Figure 3: Genetic ancestry analysis of SMC donor population.** Principal component analysis of the genotypes of the 151 donors in our population and 1000 Genomes populations. The colors indicate different 1000 Genomes reference samples. Donors are represented with "+" and black color.

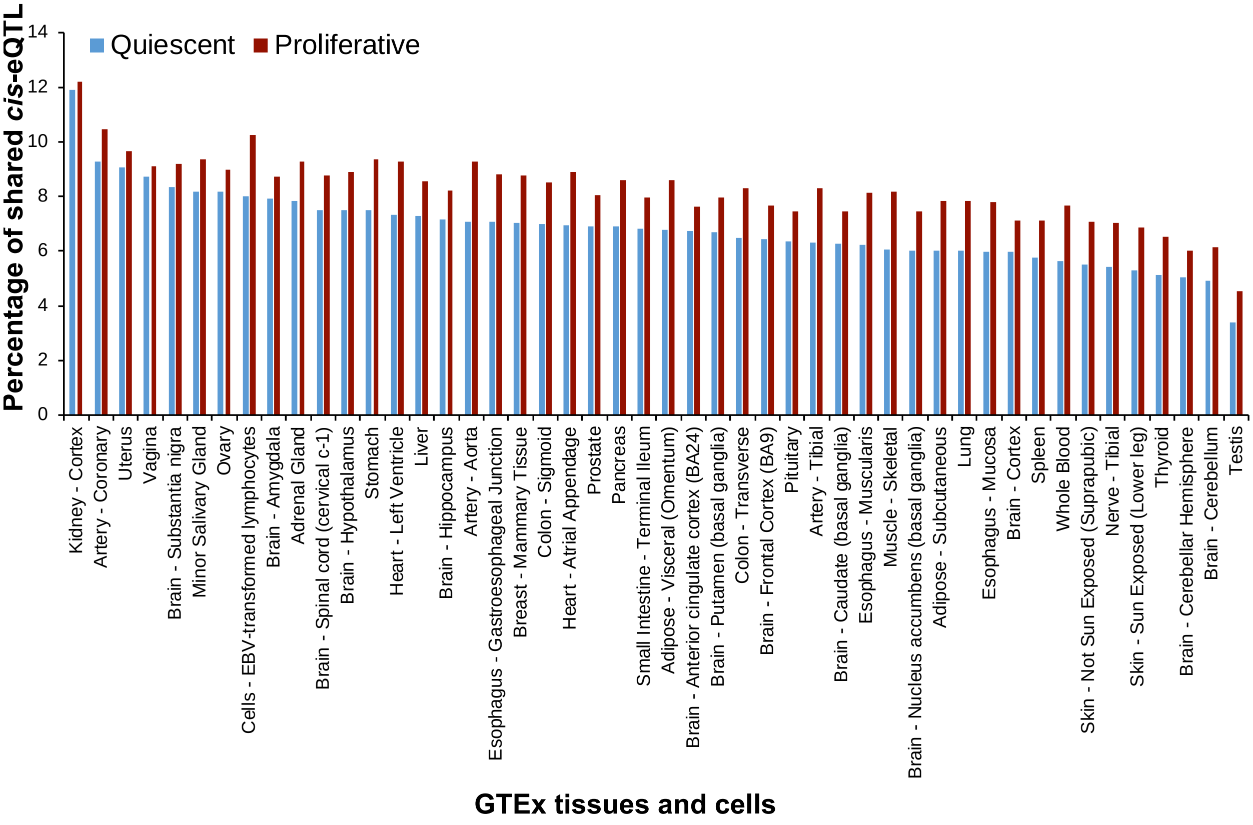


**Supplementary Figure 4: SMC-specific eQTLs.** SMC-specific eQTLs were identified using METASOFT[48]. 29 SMC-specific eQTLs were defined by SNP-gene pairs with posterior probability (METASOFT M-value) > 0.9 in SMCs and < 0.1 in all the GTEx tissues and cells. The plots show the comparison of the effect sizes of SMC-specific eQTLs and 49 GTEx tissues and cells. Error bar indicates 95% confidence intervals.

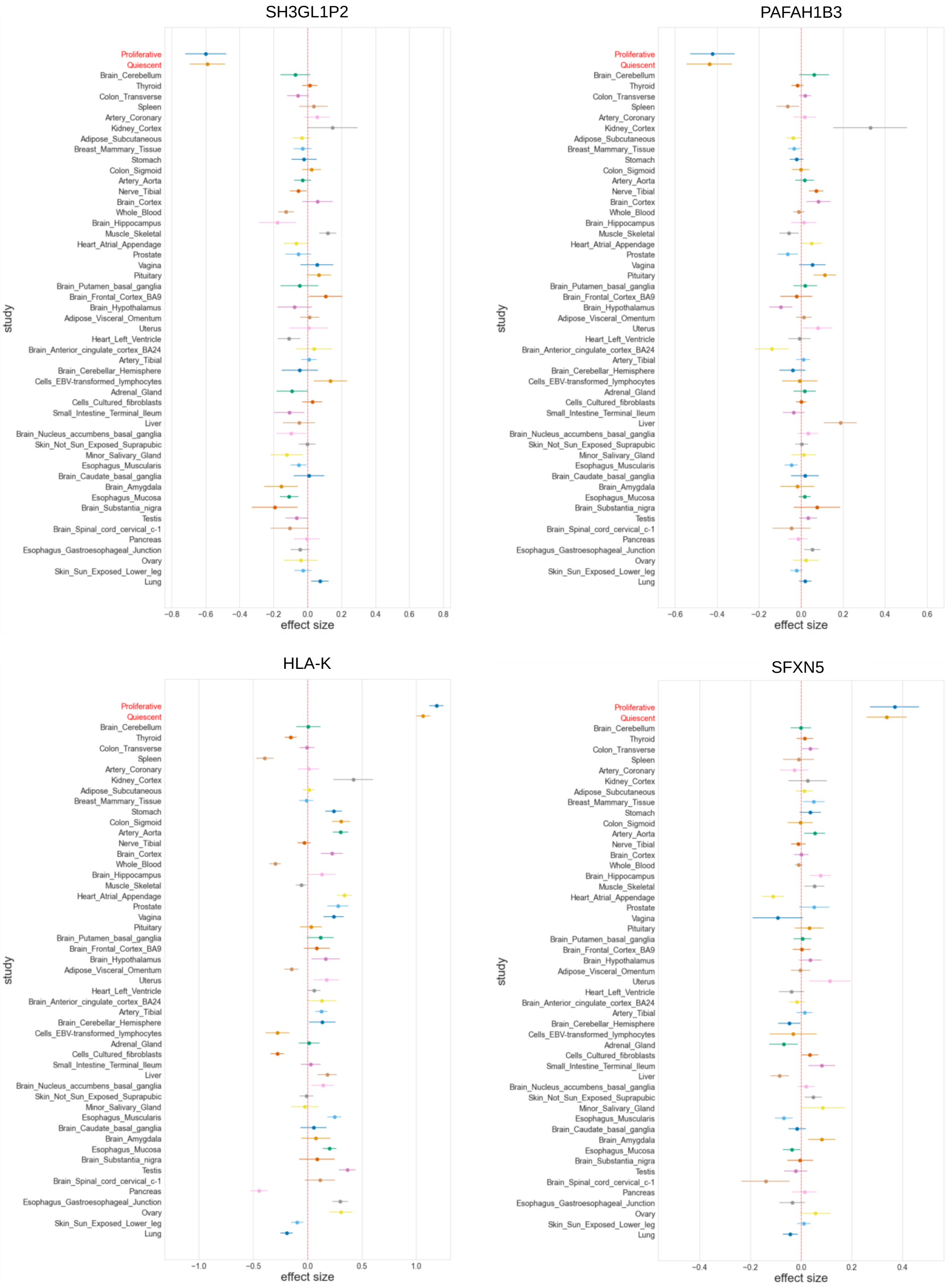
SH3GL1P2
PAFAH1B3
HLA-K
SFXN5
study
effect size
Proliferative
Quiescent
Brain_Cerebellum
Thyroid
Colon_Transverse
Spleen
Artery_Coronary
Kidney_Cortex
Adipose_Subcutaneous
Breast_Mammary_Tissue
Stomach
Colon_Sigmoid
Artery_Aorta
Nerve_Tibial
Brain_Cortex
Whole_Blood
Brain_Hippocampus
Muscle_Skeletal
Heart_Atrial_Appendage
Prostate
Vagina
Pituitary
Brain_Putamen_basal_ganglia
Brain_Frontal_Cortex_BA9
Brain_Hypothalamus
Adipose_Visceral_Omentum
Uterus
Heart_Left_Ventricle
Brain_Anterior_cingulate_cortex_BA24
Artery_Tibial
Brain_Cerebellar_Hemisphere
Cells_EBV-transformed_lymphocytes
Adrenal_Gland
Cells_Cultured_fibroblasts
Small_Intestine_Terminal_Ileum
Liver
Brain_Nucleus_accumbens_basal_ganglia
Skin_Not_Sun_Exposed_Suprapubic
Minor_Salivary_Gland
Esophagus_Muscularis
Brain_Caudate_basal_ganglia
Brain_Amygdala
Esophagus_Mucosa
Brain_Substantia_nigra
Testis
Brain_Spinal_cord_cervical_c-1
Pancreas
Esophagus_Gastroesophageal_Junction
Ovary
Skin_Sun_Exposed_Lower_leg
Lung
−0.8 −0.6 −0.4 −0.2 0.0 0.2 0.4 0.6 0.8
−0.6 −0.4 −0.2 0.0 0.2 0.4 0.6
−1.0 −0.5 0.0 0.5 1.0
−0.4 −0.2 0.0 0.2 0.4

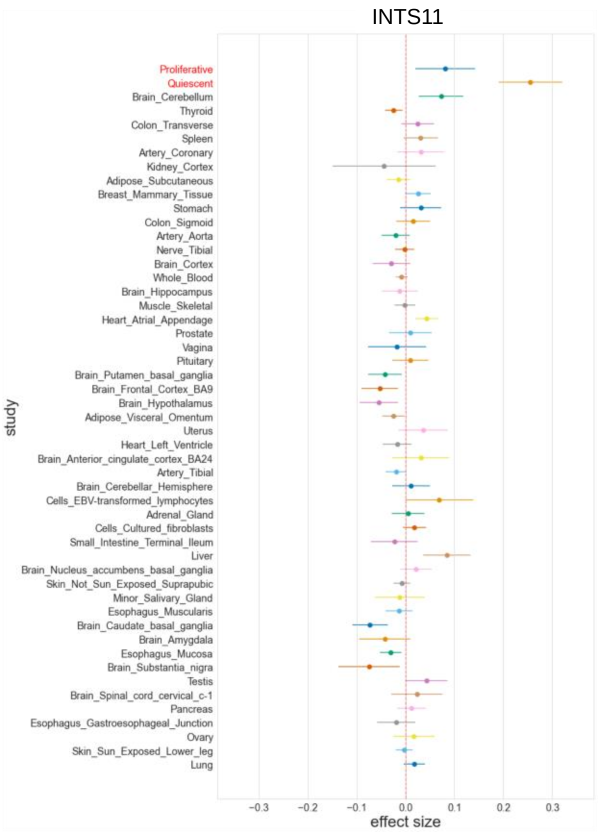
INTS11
Proliferative
Quiescent
Brain_Cerebellum
Thyroid
Colon_Transverse
Spleen
Artery_Coronary
Kidney_Cortex
Adipose_Subcutaneous
Breast_Mammary_Tissue
Stomach
Colon_Sigmoid
Artery_Aorta
Nerve_Tibial
Brain_Cortex
Whole_Blood
Brain_Hippocampus
Muscle_Skeletal
Heart_Atrial_Appendage
Prostate
Vagina
Pituitary
Brain_Putamen_basal_ganglia
Brain_Frontal_Cortex_BA9
Brain_Hypothalamus
Adipose_Visceral_Omentum
Uterus
Heart_Left_Ventricle
Brain_Anterior_cingulate_cortex_BA24
Artery_Tibial
Brain_Cerebellar_Hemisphere
Cells_EBV-transformed_lymphocytes
Adrenal_Gland
Cells_Cultured_fibroblasts
Small_Intestine_Terminal_Ileum
Liver
Brain_Nucleus_accumbens_basal_ganglia
Skin_Not_Sun_Exposed_Suprapubic
Minor_Salivary_Gland
Esophagus_Muscularis
Brain_Caudate_basal_ganglia
Brain_Amygdala
Esophagus_Mucosa
Brain_Substantia_nigra
Testis
Brain_Spinal_cord_cervical_c-1
Pancreas
Esophagus_Gastroesophageal_Junction
Ovary
Skin_Sun_Exposed_Lower_leg
Lung
study
-0.3
-0.2
-0.1
0.0
0.1
0.2
0.3
effect size

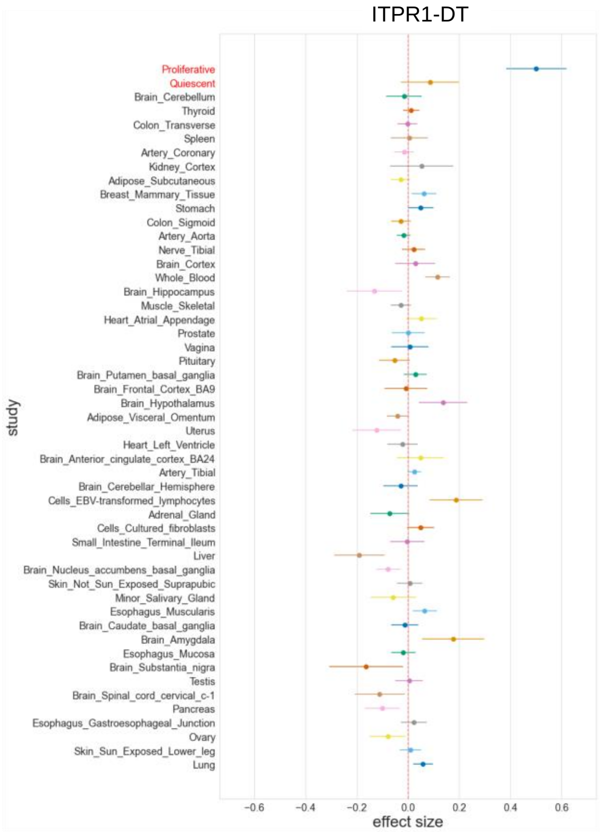
ITPR1-DT
Proliferative
Quiescent
Brain_Cerebellum
Thyroid
Colon_Transverse
Spleen
Artery_Coronary
Kidney_Cortex
Adipose_Subcutaneous
Breast_Mammary_Tissue
Stomach
Colon_Sigmoid
Artery_Aorta
Nerve_Tibial
Brain_Cortex
Whole_Blood
Brain_Hippocampus
Muscle_Skeletal
Heart_Atrial_Appendage
Prostate
Vagina
Pituitary
Brain_Putamen_basal_ganglia
Brain_Frontal_Cortex_BA9
Brain_Hypothalamus
Adipose_Visceral_Omentum
Uterus
Heart_Left_Ventricle
Brain_Anterior_cingulate_cortex_BA24
Artery_Tibial
Brain_Cerebellar_Hemisphere
Cells_EBV-transformed_lymphocytes
Adrenal_Gland
Cells_Cultured_fibroblasts
Small_Intestine_Terminal_Ileum
Liver
Brain_Nucleus_accumbens_basal_ganglia
Skin_Not_Sun_Exposed_Suprapubic
Minor_Salivary_Gland
Esophagus_Muscularis
Brain_Caudate_basal_ganglia
Brain_Amygdala
Esophagus_Mucosa
Brain_Substantia_nigra
Testis
Brain_Spinal_cord_cervical_c-1
Pancreas
Esophagus_Gastroesophageal_Junction
Ovary
Skin_Sun_Exposed_Lower_leg
Lung
study
-0.6
-0.4
-0.2
0.0
0.2
0.4
0.6
effect size

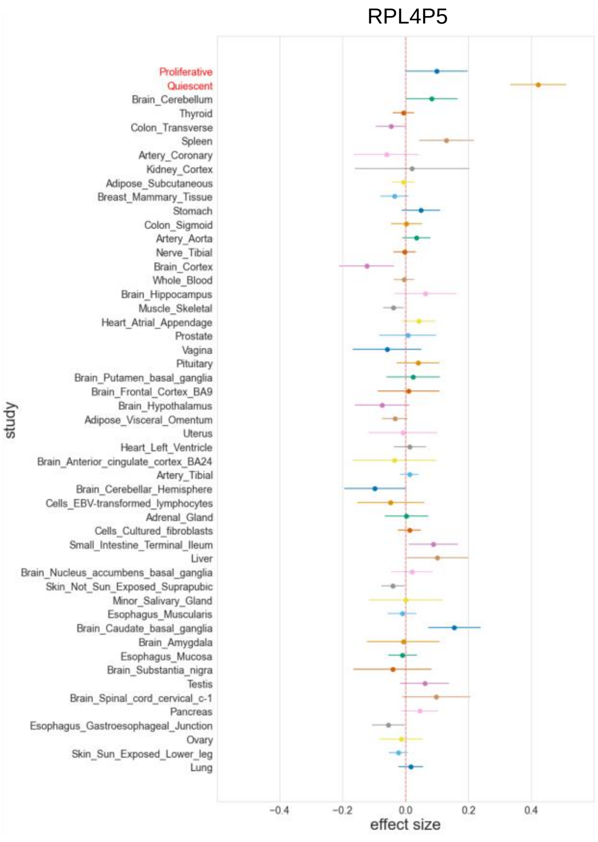
RPL4P5
Proliferative
Quiescent
Brain_Cerebellum
Thyroid
Colon_Transverse
Spleen
Artery_Coronary
Kidney_Cortex
Adipose_Subcutaneous
Breast_Mammary_Tissue
Stomach
Colon_Sigmoid
Artery_Aorta
Nerve_Tibial
Brain_Cortex
Whole_Blood
Brain_Hippocampus
Muscle_Skeletal
Heart_Atrial_Appendage
Prostate
Vagina
Pituitary
Brain_Putamen_basal_ganglia
Brain_Frontal_Cortex_BA9
Brain_Hypothalamus
Adipose_Visceral_Omentum
Uterus
Heart_Left_Ventricle
Brain_Anterior_cingulate_cortex_BA24
Artery_Tibial
Brain_Cerebellar_Hemisphere
Cells_EBV-transformed_lymphocytes
Adrenal_Gland
Cells_Cultured_fibroblasts
Small_Intestine_Terminal_Ileum
Liver
Brain_Nucleus_accumbens_basal_ganglia
Skin_Not_Sun_Exposed_Suprapubic
Minor_Salivary_Gland
Esophagus_Muscularis
Brain_Caudate_basal_ganglia
Brain_Amygdala
Esophagus_Mucosa
Brain_Substantia_nigra
Testis
Brain_Spinal_cord_cervical_c-1
Pancreas
Esophagus_Gastroesophageal_Junction
Ovary
Skin_Sun_Exposed_Lower_leg
Lung
study
-0.4
-0.2
0.0
0.2
0.4
effect size

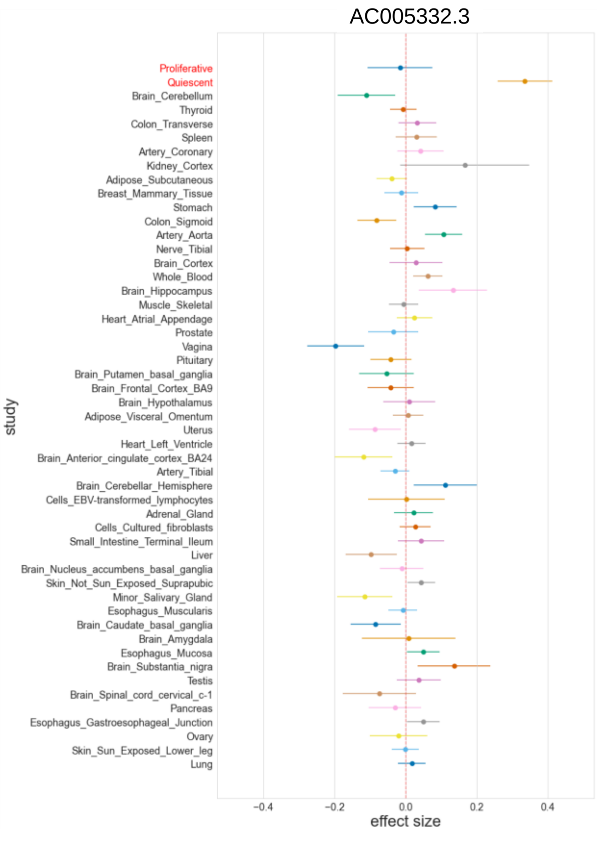
AC005332.3
Proliferative
Quiescent
Brain_Cerebellum
Thyroid
Colon_Transverse
Spleen
Artery_Coronary
Kidney_Cortex
Adipose_Subcutaneous
Breast_Mammary_Tissue
Stomach
Colon_Sigmoid
Artery_Aorta
Nerve_Tibial
Brain_Cortex
Whole_Blood
Brain_Hippocampus
Muscle_Skeletal
Heart_Atrial_Appendage
Prostate
Vagina
Pituitary
Brain_Putamen_basal_ganglia
Brain_Frontal_Cortex_BA9
Brain_Hypothalamus
Adipose_Visceral_Omentum
Uterus
Heart_Left_Ventricle
Brain_Anterior_cingulate_cortex_BA24
Artery_Tibial
Brain_Cerebellar_Hemisphere
Cells_EBV-transformed_lymphocytes
Adrenal_Gland
Cells_Cultured_fibroblasts
Small_Intestine_Terminal_Ileum
Liver
Brain_Nucleus_accumbens_basal_ganglia
Skin_Not_Sun_Exposed_Suprapubic
Minor_Salivary_Gland
Esophagus_Muscularis
Brain_Caudate_basal_ganglia
Brain_Amygdala
Esophagus_Mucosa
Brain_Substantia_nigra
Testis
Brain_Spinal_cord_cervical_c-1
Pancreas
Esophagus_Gastroesophageal_Junction
Ovary
Skin_Sun_Exposed_Lower_leg
Lung
study
-0.4
-0.2
0.0
0.2
0.4
effect size

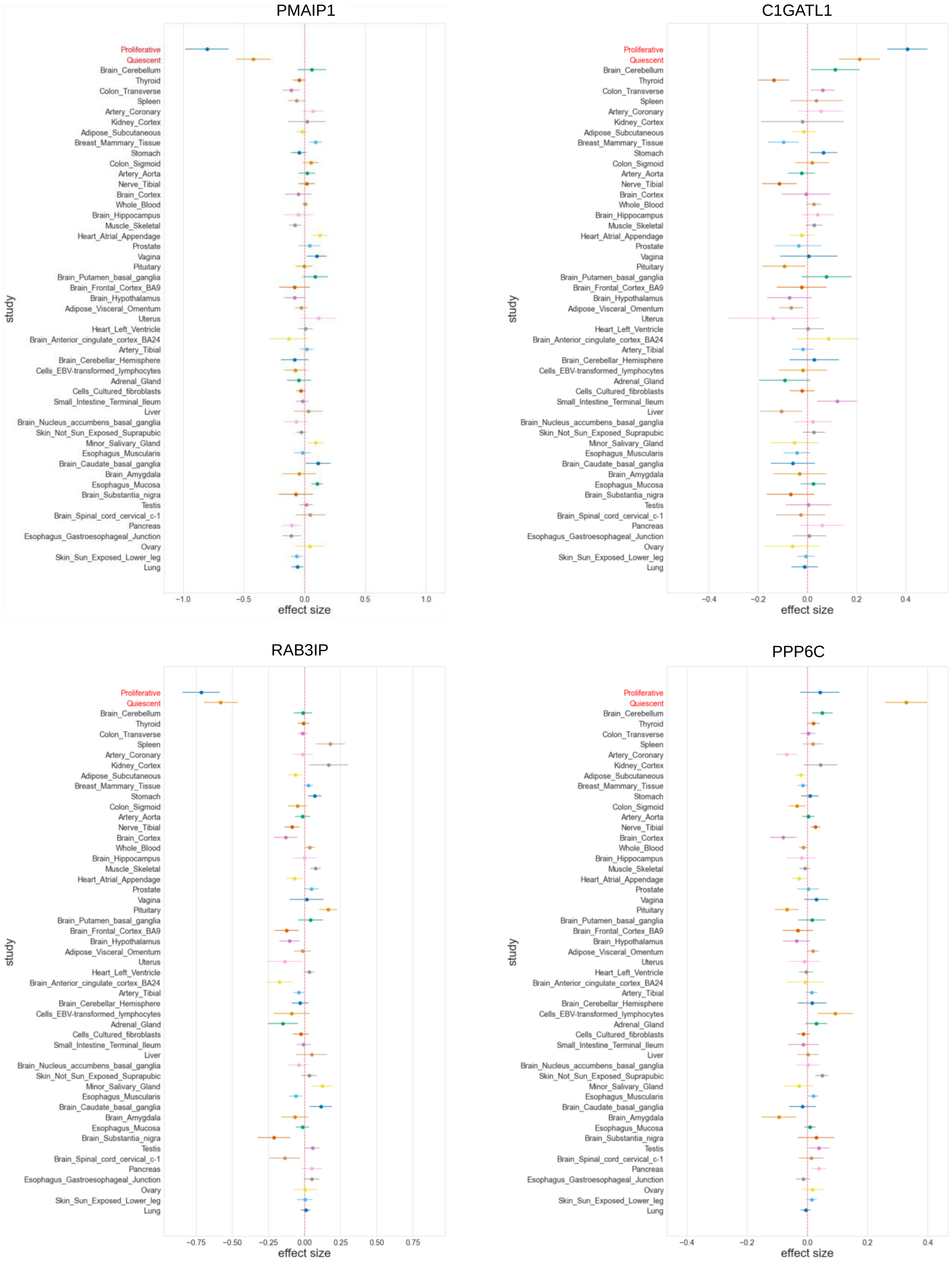
PMAIP1
C1GATL1
RAB3IP
PPP6C
study
Proliferative
Quiescent
Brain_Cerebellum
Thyroid
Colon_Transverse
Spleen
Artery_Coronary
Kidney_Cortex
Adipose_Subcutaneous
Breast_Mammary_Tissue
Stomach
Colon_Sigmoid
Artery_Aorta
Nerve_Tibial
Brain_Cortex
Whole_Blood
Brain_Hippocampus
Muscle_Skeletal
Heart_Atrial_Appendage
Prostate
Vagina
Pituitary
Brain_Putamen_basal_ganglia
Brain_Frontal_Cortex_BA9
Brain_Hypothalamus
Adipose_Visceral_Omentum
Uterus
Heart_Left_Ventricle
Brain_Anterior_cingulate_cortex_BA24
Artery_Tibial
Brain_Cerebellar_Hemisphere
Cells_EBV-transformed_lymphocytes
Adrenal_Gland
Cells_Cultured_fibroblasts
Small_Intestine_Terminal_Ileum
Liver
Brain_Nucleus_accumbens_basal_ganglia
Skin_Not_Sun_Exposed_Suprapubic
Minor_Salivary_Gland
Esophagus_Muscularis
Brain_Caudate_basal_ganglia
Brain_Amygdala
Esophagus_Mucosa
Brain_Substantia_nigra
Testis
Brain_Spinal_cord_cervical_c-1
Pancreas
Esophagus_Gastroesophageal_Junction
Ovary
Skin_Sun_Exposed_Lower_leg
Lung
effect size
−1.0 −0.5 0.0 0.5 1.0
−0.4 −0.2 0.0 0.2 0.4
−0.75 −0.50 −0.25 0.00 0.25 0.50 0.75
−0.4 −0.2 0.0 0.2 0.4

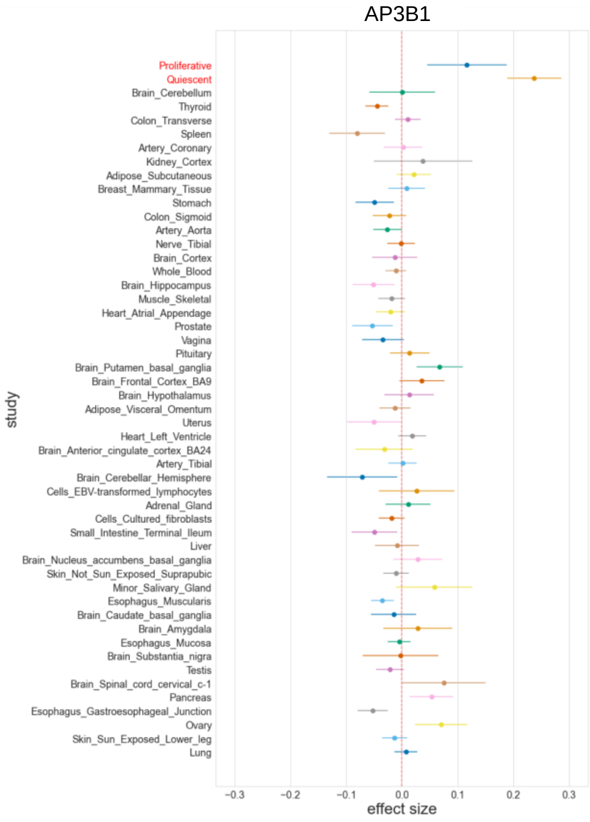
AP3B1
Proliferative
Quiescent
Brain_Cerebellum
Thyroid
Colon_Transverse
Spleen
Artery_Coronary
Kidney_Cortex
Adipose_Subcutaneous
Breast_Mammary_Tissue
Stomach
Colon_Sigmoid
Artery_Aorta
Nerve_Tibial
Brain_Cortex
Whole_Blood
Brain_Hippocampus
Muscle_Skeletal
Heart_Atrial_Appendage
Prostate
Vagina
Pituitary
Brain_Putamen_basal_ganglia
Brain_Frontal_Cortex_BA9
Brain_Hypothalamus
Adipose_Visceral_Omentum
Uterus
Heart_Left_Ventricle
Brain_Anterior_cingulate_cortex_BA24
Artery_Tibial
Brain_Cerebellar_Hemisphere
Cells_EBV-transformed_lymphocytes
Adrenal_Gland
Cells_Cultured_fibroblasts
Small_Intestine_Terminal_Ileum
Liver
Brain_Nucleus_accumbens_basal_ganglia
Skin_Not_Sun_Exposed_Suprapubic
Minor_Salivary_Gland
Esophagus_Muscularis
Brain_Caudate_basal_ganglia
Brain_Amygdala
Esophagus_Mucosa
Brain_Substantia_nigra
Testis
Brain_Spinal_cord_cervical_c-1
Pancreas
Esophagus_Gastroesophageal_Junction
Ovary
Skin_Sun_Exposed_Lower_leg
Lung
study
−0.3 −0.2 −0.1 0.0 0.1 0.2 0.3
effect size

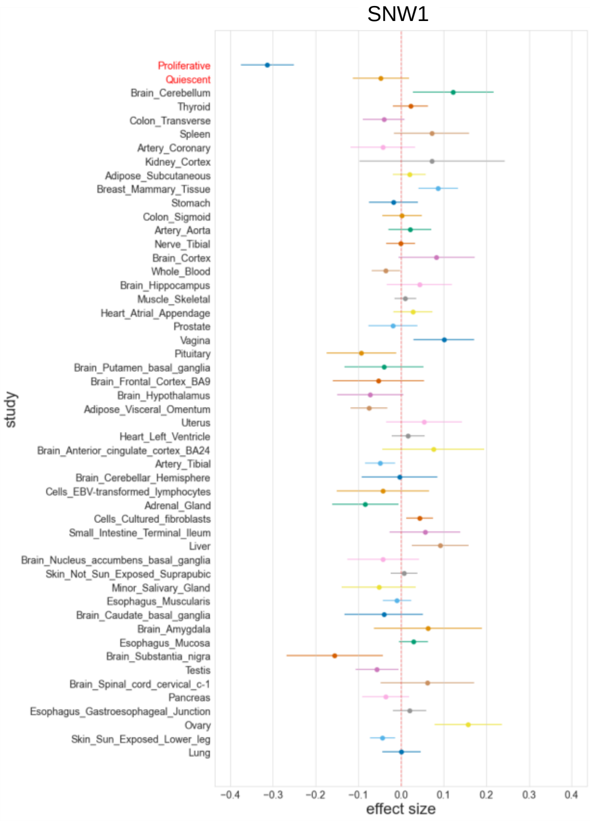
SNW1
Proliferative
Quiescent
Brain_Cerebellum
Thyroid
Colon_Transverse
Spleen
Artery_Coronary
Kidney_Cortex
Adipose_Subcutaneous
Breast_Mammary_Tissue
Stomach
Colon_Sigmoid
Artery_Aorta
Nerve_Tibial
Brain_Cortex
Whole_Blood
Brain_Hippocampus
Muscle_Skeletal
Heart_Atrial_Appendage
Prostate
Vagina
Pituitary
Brain_Putamen_basal_ganglia
Brain_Frontal_Cortex_BA9
Brain_Hypothalamus
Adipose_Visceral_Omentum
Uterus
Heart_Left_Ventricle
Brain_Anterior_cingulate_cortex_BA24
Artery_Tibial
Brain_Cerebellar_Hemisphere
Cells_EBV-transformed_lymphocytes
Adrenal_Gland
Cells_Cultured_fibroblasts
Small_Intestine_Terminal_Ileum
Liver
Brain_Nucleus_accumbens_basal_ganglia
Skin_Not_Sun_Exposed_Suprapubic
Minor_Salivary_Gland
Esophagus_Muscularis
Brain_Caudate_basal_ganglia
Brain_Amygdala
Esophagus_Mucosa
Brain_Substantia_nigra
Testis
Brain_Spinal_cord_cervical_c-1
Pancreas
Esophagus_Gastroesophageal_Junction
Ovary
Skin_Sun_Exposed_Lower_leg
Lung
study
−0.4 −0.3 −0.2 −0.1 0.0 0.1 0.2 0.3 0.4
effect size

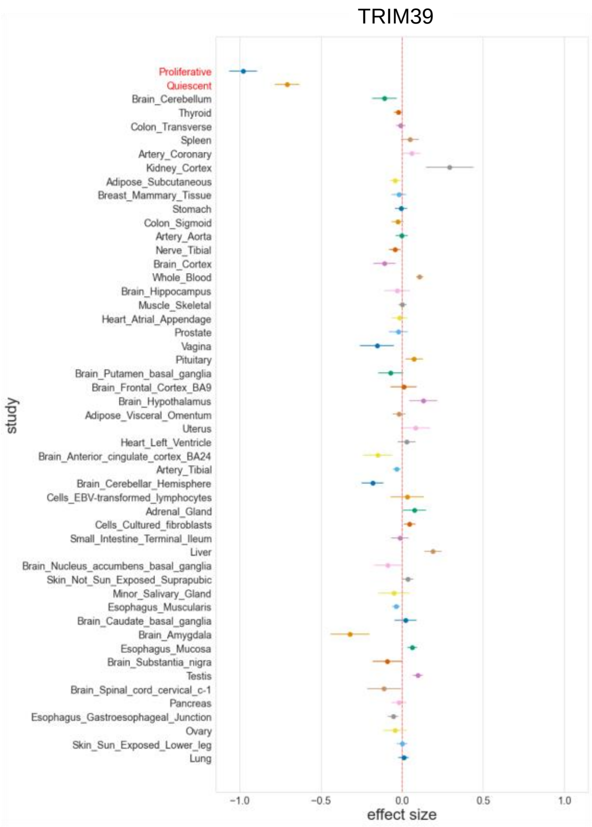
TRIM39
Proliferative
Quiescent
Brain_Cerebellum
Thyroid
Colon_Transverse
Spleen
Artery_Coronary
Kidney_Cortex
Adipose_Subcutaneous
Breast_Mammary_Tissue
Stomach
Colon_Sigmoid
Artery_Aorta
Nerve_Tibial
Brain_Cortex
Whole_Blood
Brain_Hippocampus
Muscle_Skeletal
Heart_Atrial_Appendage
Prostate
Vagina
Pituitary
Brain_Putamen_basal_ganglia
Brain_Frontal_Cortex_BA9
Brain_Hypothalamus
Adipose_Visceral_Omentum
Uterus
Heart_Left_Ventricle
Brain_Anterior_cingulate_cortex_BA24
Artery_Tibial
Brain_Cerebellar_Hemisphere
Cells_EBV-transformed_lymphocytes
Adrenal_Gland
Cells_Cultured_fibroblasts
Small_Intestine_Terminal_Ileum
Liver
Brain_Nucleus_accumbens_basal_ganglia
Skin_Not_Sun_Exposed_Suprapubic
Minor_Salivary_Gland
Esophagus_Muscularis
Brain_Caudate_basal_ganglia
Brain_Amygdala
Esophagus_Mucosa
Brain_Substantia_nigra
Testis
Brain_Spinal_cord_cervical_c-1
Pancreas
Esophagus_Gastroesophageal_Junction
Ovary
Skin_Sun_Exposed_Lower_leg
Lung
study
−1.0 −0.5 0.0 0.5 1.0
effect size

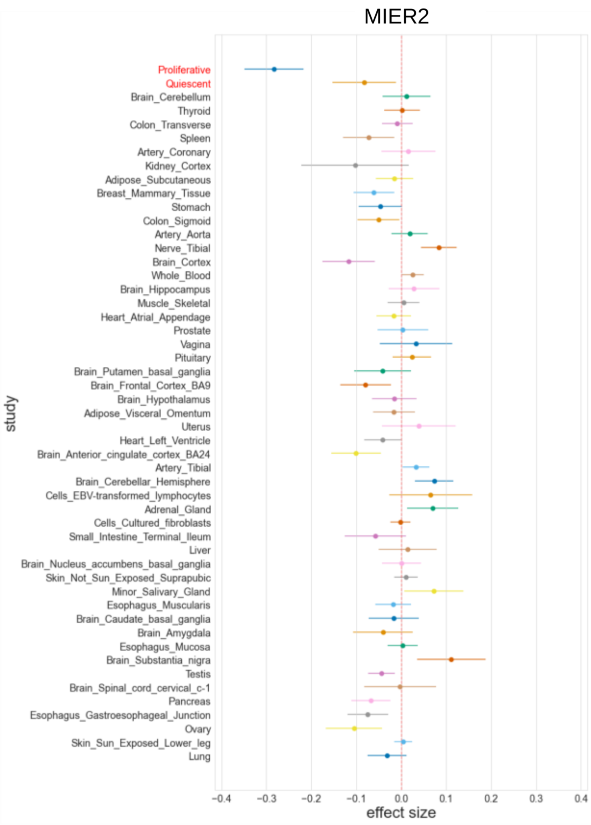
MIER2
Proliferative
Quiescent
Brain_Cerebellum
Thyroid
Colon_Transverse
Spleen
Artery_Coronary
Kidney_Cortex
Adipose_Subcutaneous
Breast_Mammary_Tissue
Stomach
Colon_Sigmoid
Artery_Aorta
Nerve_Tibial
Brain_Cortex
Whole_Blood
Brain_Hippocampus
Muscle_Skeletal
Heart_Atrial_Appendage
Prostate
Vagina
Pituitary
Brain_Putamen_basal_ganglia
Brain_Frontal_Cortex_BA9
Brain_Hypothalamus
Adipose_Visceral_Omentum
Uterus
Heart_Left_Ventricle
Brain_Anterior_cingulate_cortex_BA24
Artery_Tibial
Brain_Cerebellar_Hemisphere
Cells_EBV-transformed_lymphocytes
Adrenal_Gland
Cells_Cultured_fibroblasts
Small_Intestine_Terminal_Ileum
Liver
Brain_Nucleus_accumbens_basal_ganglia
Skin_Not_Sun_Exposed_Suprapubic
Minor_Salivary_Gland
Esophagus_Muscularis
Brain_Caudate_basal_ganglia
Brain_Amygdala
Esophagus_Mucosa
Brain_Substantia_nigra
Testis
Brain_Spinal_cord_cervical_c-1
Pancreas
Esophagus_Gastroesophageal_Junction
Ovary
Skin_Sun_Exposed_Lower_leg
Lung
study
−0.4 −0.3 −0.2 −0.1 0.0 0.1 0.2 0.3 0.4
effect size

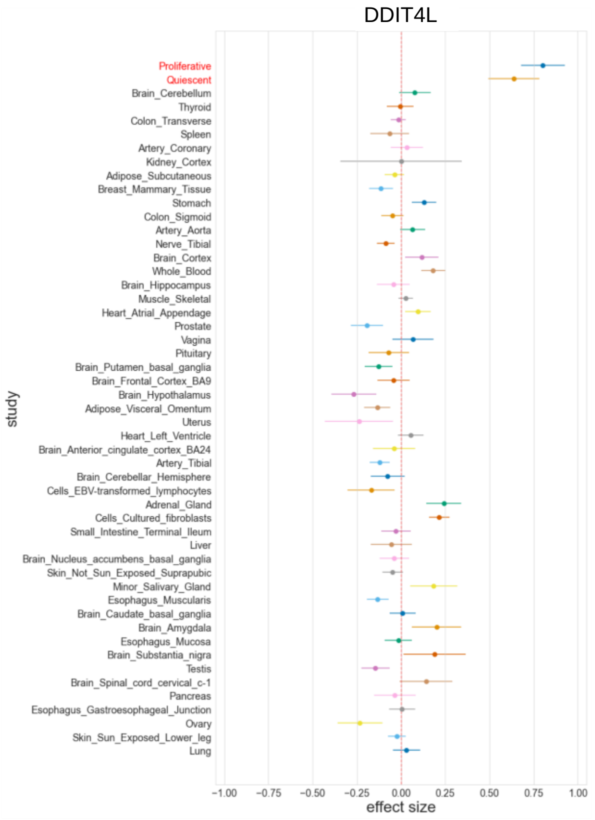
DDIT4L
Proliferative
Quiescent
Brain_Cerebellum
Thyroid
Colon_Transverse
Spleen
Artery_Coronary
Kidney_Cortex
Adipose_Subcutaneous
Breast_Mammary_Tissue
Stomach
Colon_Sigmoid
Artery_Aorta
Nerve_Tibial
Brain_Cortex
Whole_Blood
Brain_Hippocampus
Muscle_Skeletal
Heart_Atrial_Appendage
Prostate
Vagina
Pituitary
Brain_Putamen_basal_ganglia
Brain_Frontal_Cortex_BA9
Brain_Hypothalamus
Adipose_Visceral_Omentum
Uterus
Heart_Left_Ventricle
Brain_Anterior_cingulate_cortex_BA24
Artery_Tibial
Brain_Cerebellar_Hemisphere
Cells_EBV-transformed_lymphocytes
Adrenal_Gland
Cells_Cultured_fibroblasts
Small_Intestine_Terminal_Ileum
Liver
Brain_Nucleus_accumbens_basal_ganglia
Skin_Not_Sun_Exposed_Suprapubic
Minor_Salivary_Gland
Esophagus_Muscularis
Brain_Caudate_basal_ganglia
Brain_Amygdala
Esophagus_Mucosa
Brain_Substantia_nigra
Testis
Brain_Spinal_cord_cervical_c-1
Pancreas
Esophagus_Gastroesophageal_Junction
Ovary
Skin_Sun_Exposed_Lower_leg
Lung
study
−1.00 −0.75 −0.50 −0.25 0.00 0.25 0.50 0.75 1.00
effect size

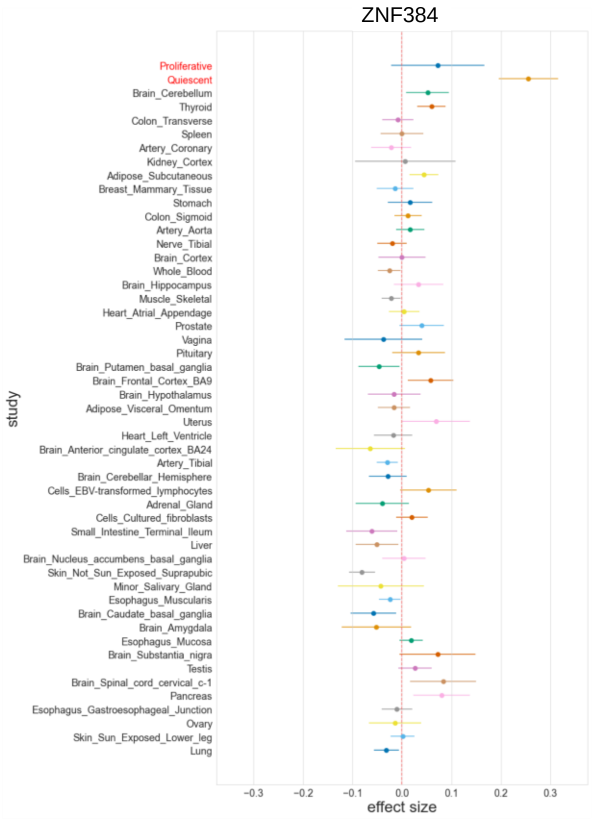
ZNF384
Proliferative
Quiescent
Brain_Cerebellum
Thyroid
Colon_Transverse
Spleen
Artery_Coronary
Kidney_Cortex
Adipose_Subcutaneous
Breast_Mammary_Tissue
Stomach
Colon_Sigmoid
Artery_Aorta
Nerve_Tibial
Brain_Cortex
Whole_Blood
Brain_Hippocampus
Muscle_Skeletal
Heart_Atrial_Appendage
Prostate
Vagina
Pituitary
Brain_Putamen_basal_ganglia
Brain_Frontal_Cortex_BA9
Brain_Hypothalamus
Adipose_Visceral_Omentum
Uterus
Heart_Left_Ventricle
Brain_Anterior_cingulate_cortex_BA24
Artery_Tibial
Brain_Cerebellar_Hemisphere
Cells_EBV-transformed_lymphocytes
Adrenal_Gland
Cells_Cultured_fibroblasts
Small_Intestine_Terminal_Ileum
Liver
Brain_Nucleus_accumbens_basal_ganglia
Skin_Not_Sun_Exposed_Suprapubic
Minor_Salivary_Gland
Esophagus_Muscularis
Brain_Caudate_basal_ganglia
Brain_Amygdala
Esophagus_Mucosa
Brain_Substantia_nigra
Testis
Brain_Spinal_cord_cervical_c-1
Pancreas
Esophagus_Gastroesophageal_Junction
Ovary
Skin_Sun_Exposed_Lower_leg
Lung
study
−0.3 −0.2 −0.1 0.0 0.1 0.2 0.3
effect size

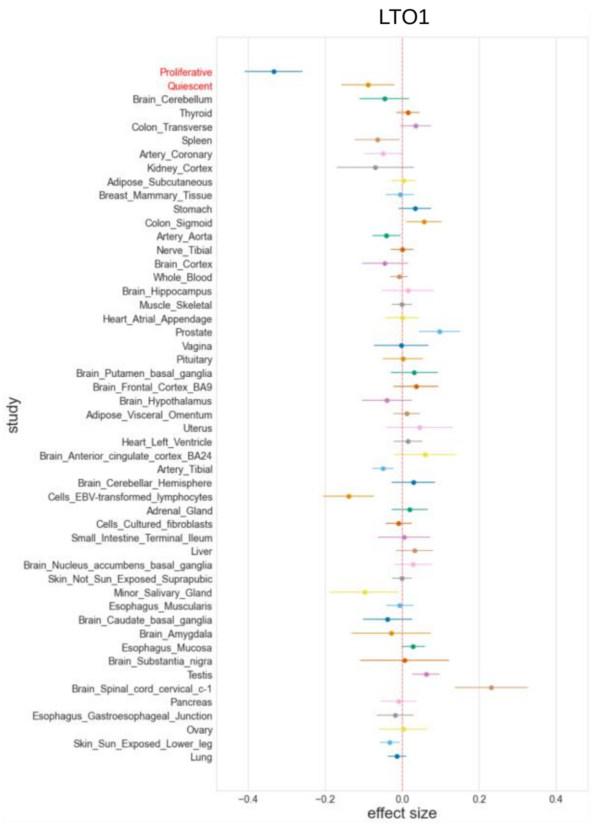
LTO1
Proliferative
Quiescent
Brain_Cerebellum
Thyroid
Colon_Transverse
Spleen
Artery_Coronary
Kidney_Cortex
Adipose_Subcutaneous
Breast_Mammary_Tissue
Stomach
Colon_Sigmoid
Artery_Aorta
Nerve_Tibial
Brain_Cortex
Whole_Blood
Brain_Hippocampus
Muscle_Skeletal
Heart_Atrial_Appendage
Prostate
Vagina
Pituitary
Brain_Putamen_basal_ganglia
Brain_Frontal_Cortex_BA9
Brain_Hypothalamus
Adipose_Visceral_Omentum
Uterus
Heart_Left_Ventricle
Brain_Anterior_cingulate_cortex_BA24
Artery_Tibial
Brain_Cerebellar_Hemisphere
Cells_EBV-transformed_lymphocytes
Adrenal_Gland
Cells_Cultured_fibroblasts
Small_Intestine_Terminal_Ileum
Liver
Brain_Nucleus_accumbens_basal_ganglia
Skin_Not_Sun_Exposed_Suprapubic
Minor_Salivary_Gland
Esophagus_Muscularis
Brain_Caudate_basal_ganglia
Brain_Amygdala
Esophagus_Mucosa
Brain_Substantia_nigra
Testis
Brain_Spinal_cord_cervical_c-1
Pancreas
Esophagus_Gastroesophageal_Junction
Ovary
Skin_Sun_Exposed_Lower_leg
Lung
study
−0.4 −0.2 0.0 0.2 0.4
effect size

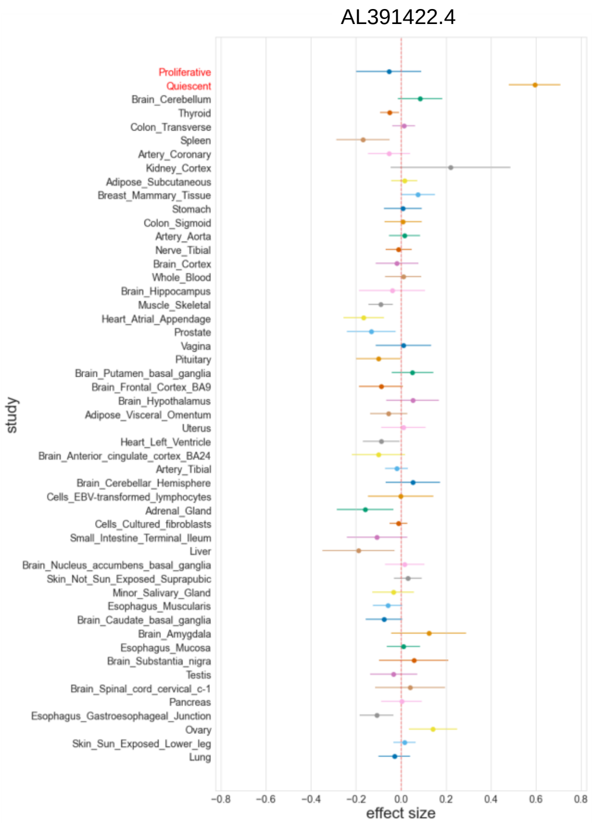
AL391422.4
Proliferative
Quiescent
Brain_Cerebellum
Thyroid
Colon_Transverse
Spleen
Artery_Coronary
Kidney_Cortex
Adipose_Subcutaneous
Breast_Mammary_Tissue
Stomach
Colon_Sigmoid
Artery_Aorta
Nerve_Tibial
Brain_Cortex
Whole_Blood
Brain_Hippocampus
Muscle_Skeletal
Heart_Atrial_Appendage
Prostate
Vagina
Pituitary
Brain_Putamen_basal_ganglia
Brain_Frontal_Cortex_BA9
Brain_Hypothalamus
Adipose_Visceral_Omentum
Uterus
Heart_Left_Ventricle
Brain_Anterior_cingulate_cortex_BA24
Artery_Tibial
Brain_Cerebellar_Hemisphere
Cells_EBV-transformed_lymphocytes
Adrenal_Gland
Cells_Cultured_fibroblasts
Small_Intestine_Terminal_Ileum
Liver
Brain_Nucleus_accumbens_basal_ganglia
Skin_Not_Sun_Exposed_Suprapubic
Minor_Salivary_Gland
Esophagus_Muscularis
Brain_Caudate_basal_ganglia
Brain_Amygdala
Esophagus_Mucosa
Brain_Substantia_nigra
Testis
Brain_Spinal_cord_cervical_c-1
Pancreas
Esophagus_Gastroesophageal_Junction
Ovary
Skin_Sun_Exposed_Lower_leg
Lung
study
−0.8 −0.6 −0.4 −0.2 0.0 0.2 0.4 0.6 0.8
effect size

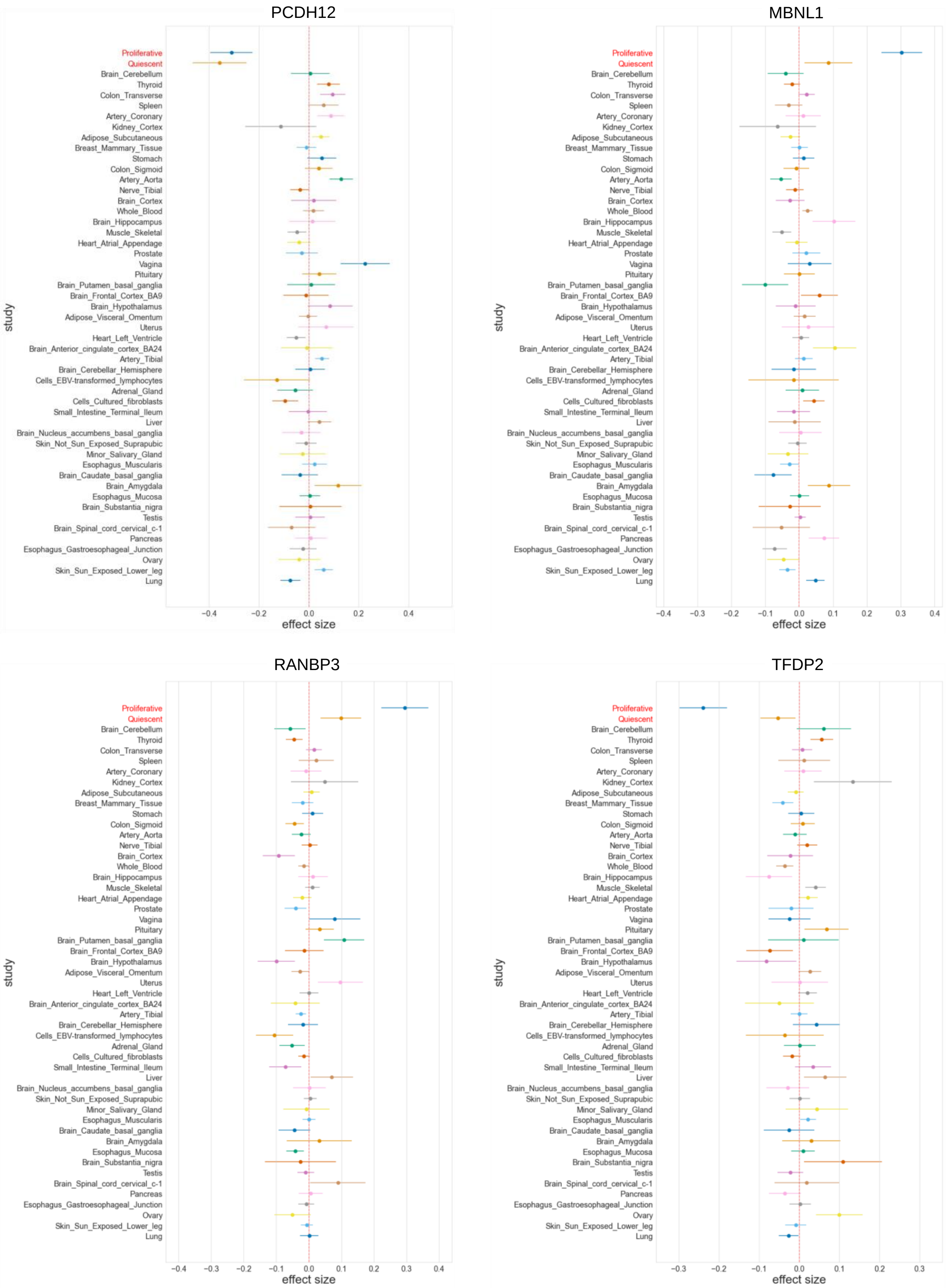
PCDH12
MBNL1
RANBP3
TFDP2
Proliferative
Quiescent
Brain_Cerebellum
Thyroid
Colon_Transverse
Spleen
Artery_Coronary
Kidney_Cortex
Adipose_Subcutaneous
Breast_Mammary_Tissue
Stomach
Colon_Sigmoid
Artery_Aorta
Nerve_Tibial
Brain_Cortex
Whole_Blood
Brain_Hippocampus
Muscle_Skeletal
Heart_Atrial_Appendage
Prostate
Vagina
Pituitary
Brain_Putamen_basal_ganglia
Brain_Frontal_Cortex_BA9
Brain_Hypothalamus
Adipose_Visceral_Omentum
Uterus
Heart_Left_Ventricle
Brain_Anterior_cingulate_cortex_BA24
Artery_Tibial
Brain_Cerebellar_Hemisphere
Cells_EBV-transformed_lymphocytes
Adrenal_Gland
Cells_Cultured_fibroblasts
Small_Intestine_Terminal_Ileum
Liver
Brain_Nucleus_accumbens_basal_ganglia
Skin_Not_Sun_Exposed_Suprapubic
Minor_Salivary_Gland
Esophagus_Muscularis
Brain_Caudate_basal_ganglia
Brain_Amygdala
Esophagus_Mucosa
Brain_Substantia_nigra
Testis
Brain_Spinal_cord_cervical_c-1
Pancreas
Esophagus_Gastroesophageal_Junction
Ovary
Skin_Sun_Exposed_Lower_leg
Lung
study
effect size

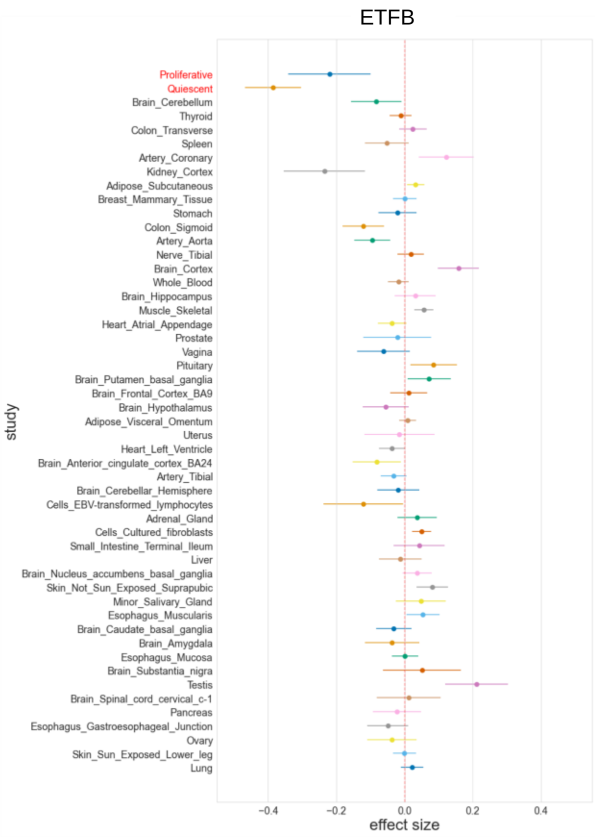
ETFB
Proliferative
Quiescent
Brain_Cerebellum
Thyroid
Colon_Transverse
Spleen
Artery_Coronary
Kidney_Cortex
Adipose_Subcutaneous
Breast_Mammary_Tissue
Stomach
Colon_Sigmoid
Artery_Aorta
Nerve_Tibial
Brain_Cortex
Whole_Blood
Brain_Hippocampus
Muscle_Skeletal
Heart_Atrial_Appendage
Prostate
Vagina
Pituitary
Brain_Putamen_basal_ganglia
Brain_Frontal_Cortex_BA9
Brain_Hypothalamus
Adipose_Visceral_Omentum
Uterus
Heart_Left_Ventricle
Brain_Anterior_cingulate_cortex_BA24
Artery_Tibial
Brain_Cerebellar_Hemisphere
Cells_EBV-transformed_lymphocytes
Adrenal_Gland
Cells_Cultured_fibroblasts
Small_Intestine_Terminal_Ileum
Liver
Brain_Nucleus_accumbens_basal_ganglia
Skin_Not_Sun_Exposed_Suprapubic
Minor_Salivary_Gland
Esophagus_Muscularis
Brain_Caudate_basal_ganglia
Brain_Amygdala
Esophagus_Mucosa
Brain_Substantia_nigra
Testis
Brain_Spinal_cord_cervical_c-1
Pancreas
Esophagus_Gastroesophageal_Junction
Ovary
Skin_Sun_Exposed_Lower_leg
Lung
study
−0.4 −0.2 0.0 0.2 0.4
effect size

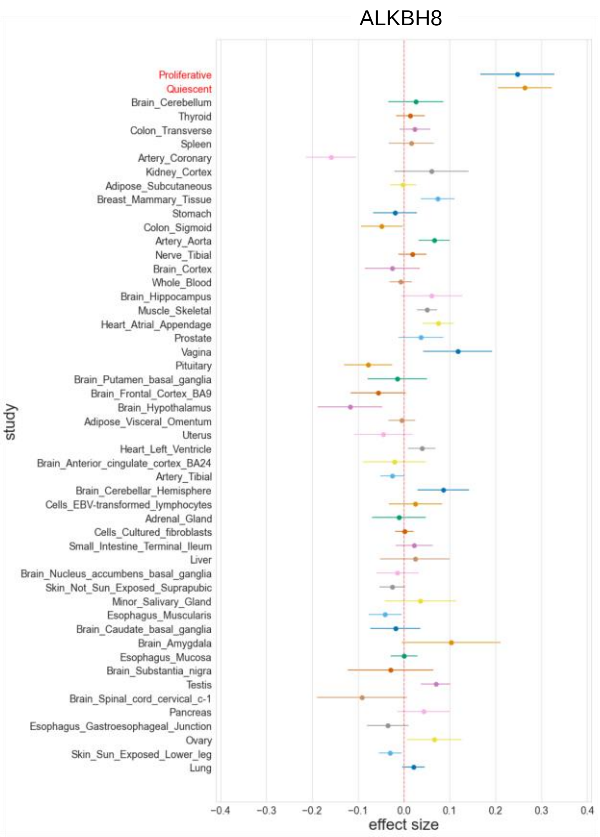
ALKBH8
Proliferative
Quiescent
Brain_Cerebellum
Thyroid
Colon_Transverse
Spleen
Artery_Coronary
Kidney_Cortex
Adipose_Subcutaneous
Breast_Mammary_Tissue
Stomach
Colon_Sigmoid
Artery_Aorta
Nerve_Tibial
Brain_Cortex
Whole_Blood
Brain_Hippocampus
Muscle_Skeletal
Heart_Atrial_Appendage
Prostate
Vagina
Pituitary
Brain_Putamen_basal_ganglia
Brain_Frontal_Cortex_BA9
Brain_Hypothalamus
Adipose_Visceral_Omentum
Uterus
Heart_Left_Ventricle
Brain_Anterior_cingulate_cortex_BA24
Artery_Tibial
Brain_Cerebellar_Hemisphere
Cells_EBV-transformed_lymphocytes
Adrenal_Gland
Cells_Cultured_fibroblasts
Small_Intestine_Terminal_Ileum
Liver
Brain_Nucleus_accumbens_basal_ganglia
Skin_Not_Sun_Exposed_Suprapubic
Minor_Salivary_Gland
Esophagus_Muscularis
Brain_Caudate_basal_ganglia
Brain_Amygdala
Esophagus_Mucosa
Brain_Substantia_nigra
Testis
Brain_Spinal_cord_cervical_c-1
Pancreas
Esophagus_Gastroesophageal_Junction
Ovary
Skin_Sun_Exposed_Lower_leg
Lung
study
−0.4 −0.3 −0.2 −0.1 0.0 0.1 0.2 0.3 0.4
effect size

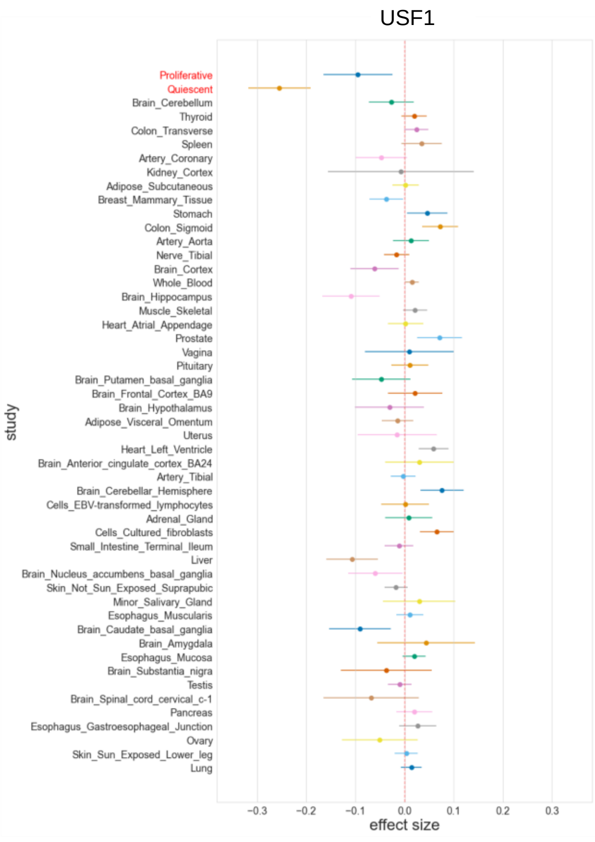
USF1
Proliferative
Quiescent
Brain_Cerebellum
Thyroid
Colon_Transverse
Spleen
Artery_Coronary
Kidney_Cortex
Adipose_Subcutaneous
Breast_Mammary_Tissue
Stomach
Colon_Sigmoid
Artery_Aorta
Nerve_Tibial
Brain_Cortex
Whole_Blood
Brain_Hippocampus
Muscle_Skeletal
Heart_Atrial_Appendage
Prostate
Vagina
Pituitary
Brain_Putamen_basal_ganglia
Brain_Frontal_Cortex_BA9
Brain_Hypothalamus
Adipose_Visceral_Omentum
Uterus
Heart_Left_Ventricle
Brain_Anterior_cingulate_cortex_BA24
Artery_Tibial
Brain_Cerebellar_Hemisphere
Cells_EBV-transformed_lymphocytes
Adrenal_Gland
Cells_Cultured_fibroblasts
Small_Intestine_Terminal_Ileum
Liver
Brain_Nucleus_accumbens_basal_ganglia
Skin_Not_Sun_Exposed_Suprapubic
Minor_Salivary_Gland
Esophagus_Muscularis
Brain_Caudate_basal_ganglia
Brain_Amygdala
Esophagus_Mucosa
Brain_Substantia_nigra
Testis
Brain_Spinal_cord_cervical_c-1
Pancreas
Esophagus_Gastroesophageal_Junction
Ovary
Skin_Sun_Exposed_Lower_leg
Lung
study
−0.3 −0.2 −0.1 0.0 0.1 0.2 0.3
effect size

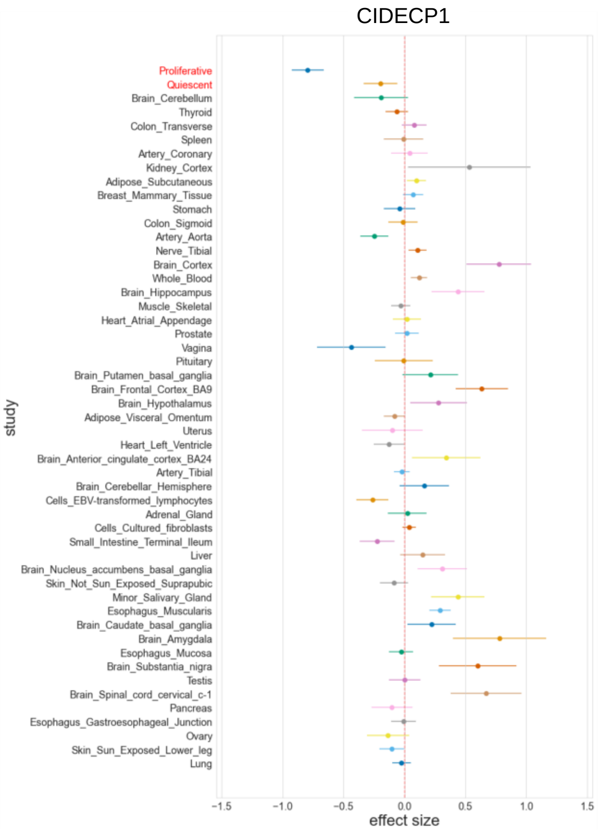
CIDECP1
Proliferative
Quiescent
Brain_Cerebellum
Thyroid
Colon_Transverse
Spleen
Artery_Coronary
Kidney_Cortex
Adipose_Subcutaneous
Breast_Mammary_Tissue
Stomach
Colon_Sigmoid
Artery_Aorta
Nerve_Tibial
Brain_Cortex
Whole_Blood
Brain_Hippocampus
Muscle_Skeletal
Heart_Atrial_Appendage
Prostate
Vagina
Pituitary
Brain_Putamen_basal_ganglia
Brain_Frontal_Cortex_BA9
Brain_Hypothalamus
Adipose_Visceral_Omentum
Uterus
Heart_Left_Ventricle
Brain_Anterior_cingulate_cortex_BA24
Artery_Tibial
Brain_Cerebellar_Hemisphere
Cells_EBV-transformed_lymphocytes
Adrenal_Gland
Cells_Cultured_fibroblasts
Small_Intestine_Terminal_Ileum
Liver
Brain_Nucleus_accumbens_basal_ganglia
Skin_Not_Sun_Exposed_Suprapubic
Minor_Salivary_Gland
Esophagus_Muscularis
Brain_Caudate_basal_ganglia
Brain_Amygdala
Esophagus_Mucosa
Brain_Substantia_nigra
Testis
Brain_Spinal_cord_cervical_c-1
Pancreas
Esophagus_Gastroesophageal_Junction
Ovary
Skin_Sun_Exposed_Lower_leg
Lung
study
−1.5 −1.0 −0.5 0.0 0.5 1.0 1.5
effect size

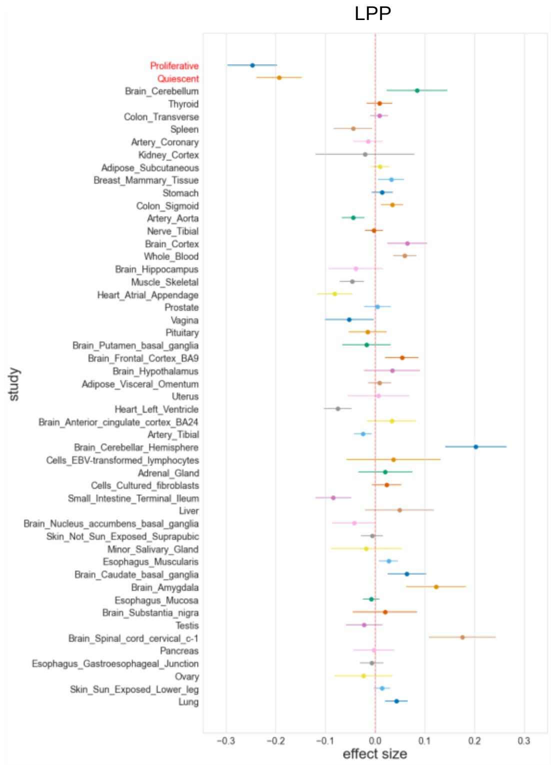


**Supplementary Figure 5: SMC-specific eQTLs.** SMC-specific eQTLs were identified using METASOFT[48]. 29 SMC-specific eQTLs were defined by SNP-gene pairs with posterior probability (METASOFT M-value) > 0.9 in SMCs and < 0.1 in all the GTEx tissues and cells. The plots show the comparison of the effect sizes of SMC-specific eQTLs and 49 GTEx tissues and cells. Error bar indicates 95% confidence intervals.

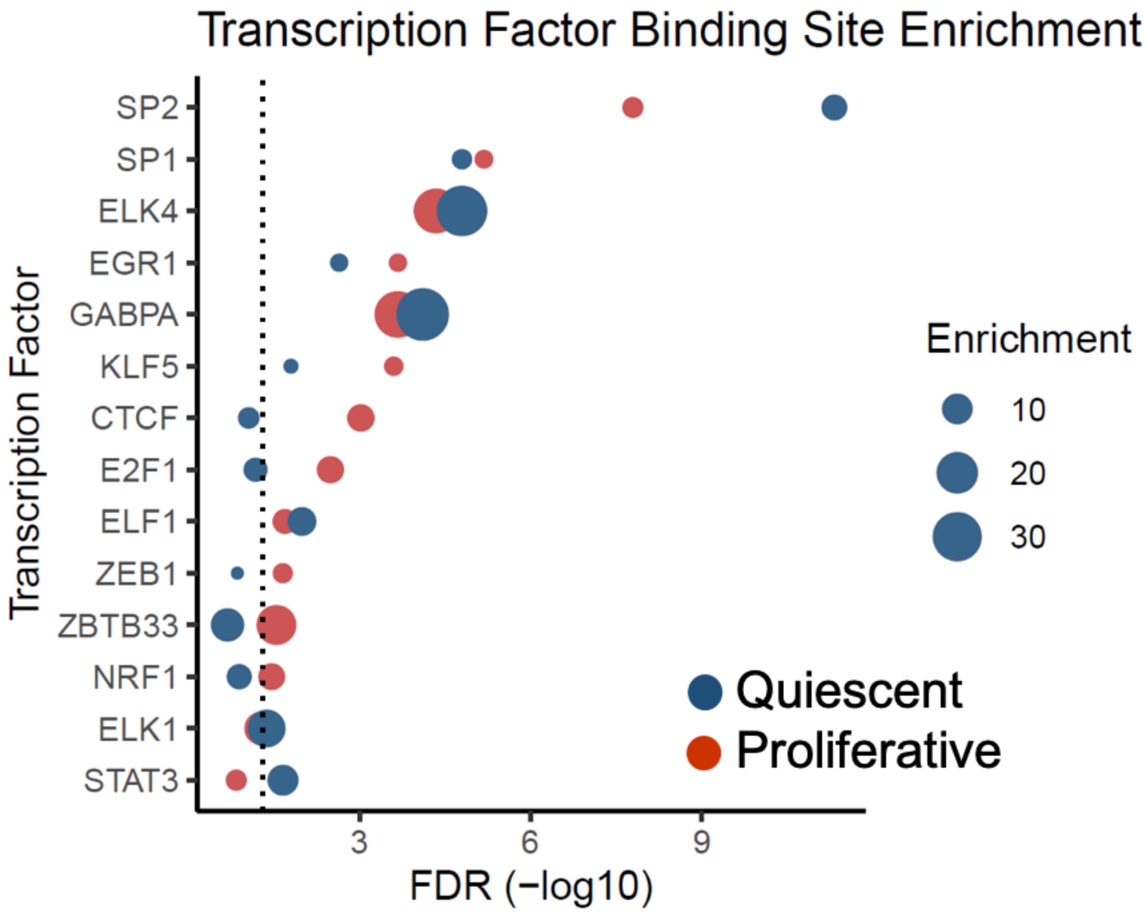


**Supplementary Figure 6: Identification of transcription factors overlapping SMC *cis*-eQTLs.** SMC *cis*-eQTL SNPs in accessible chromatin regions were interrogated for overlap for putative transcription factor (TF) binding sites using SNP2TFBS[57]. The TF enrichment values were calculated as the ratio of the observed SNP hits over the expected ones for each TF. Statistical significance of the enrichment was calculated using a binomial test and corrected for multiple testing using the qvalue package in R[40]. The plot shows TFs ranked according to their FDR q-values for both quiescent and proliferative SMCs. The point size is proportional to the TF enrichment.

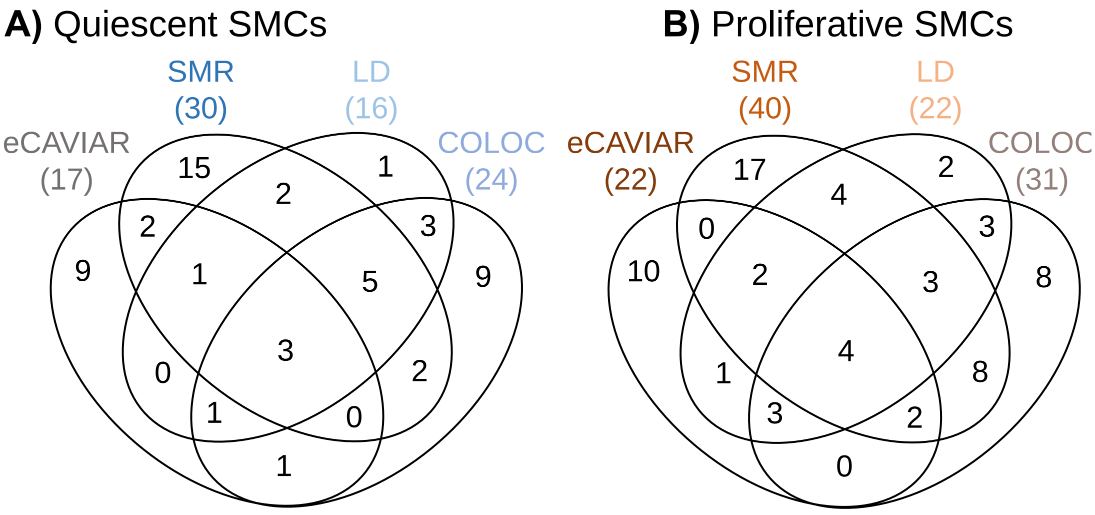


| Quiescent (17) | Both (37) | | Proliferative (30) | |
|---|---|---|---|---|
| AL513548.3$^{E,C,L,S}$ | AL592148.3$^{E}$ | LMAN1$^{E}$ | AC009054.2$^{L,S}$ | PDGFD$^{C}$ |
| CCDC158$^{C}$ | ARHGAP42$^{C,S}$ | MAP1S$^{C}$ | ADAMTS7$^{S}$ | PSD4$^{E}$ |
| HLA-B$^{S}$ | ARVCF$^{E}$ | MARCH2$^{E,C,S}$ | AP003392.4$^{E,L}$ | SKIV2L$^{S}$ |
| ILF3-AS1$^{S}$ | CFDP1$^{C,L,S}$ | MLH3$^{C,L,S}$ | ATP5MC1$^{S,L}$ | SLC2A12$^{S}$ |
| IQGAP1$^{C}$ | CGGBP1$^{E}$ | MRPL45P2$^{S}$ | CD109$^{E}$ | SMAD3$^{E,C,L,S}$ |
| LIPG$^{C}$ | DAGLB$^{C,L,S}$ | NEK9$^{C,L,S}$ | CDK8$^{E}$ | STAT6$^{C,L}$ |
| MACC1$^{E}$ | DDX59-AS1$^{C,E,L}$ | NME6$^{S}$ | CETP$^{C,L}$ | TCEA2$^{E,L,S}$ |
| PIK3CB$^{S}$ | DHODH$^{S}$ | REST$^{E,C,L,S}$ | CTSK$^{C}$ | TMEM106B$^{C}$ |
| PSMA3-AS1$^{C,L}$ | DHX38$^{C,L,S}$ | RGS19$^{L}$ | CTSS$^{S}$ | TMEM91$^{S}$ |
| RCCD1$^{C}$ | EFCAB5$^{C,S}$ | RIC8A$^{E}$ | DHX58$^{C}$ | TRIP4$^{L}$ |
| RP11-731C17.2$^{S}$ | EIF2B2$^{E,C,L,S}$ | RP11-33B1.1$^{S}$ | FURIN$^{E,C,L,S}$ | |
| RRP1B$^{E}$ | ERAP2$^{E,S}$ | SNHG18$^{C,L}$ | GOLPH3L$^{C,S}$ | |
| SERPINF1$^{S}$ | FES$^{E,C,L,S}$ | SORBS2$^{E,L,S}$ | HAUS4$^{S}$ | |
| SMG6$^{S}$ | FGD6$^{E,C,S}$ | TDRKH-AS1$^{E,C,L}$ | IRAK1BP1$^{E}$ | |
| THOC5$^{S}$ | FHL3$^{C,L,S}$ | TOM1L2$^{C,S}$ | LINC02542$^{C,S}$ | |
| TMEM87B$^{S}$ | GSTT2B$^{S}$ | ZFP30$^{S}$ | LNPEP$^{S}$ | |
| YEATS4$^{E}$ | GTF2IP12$^{C}$ | ZNF100$^{E,S}$ | MAP3K7CL$^{E,C,L,S}$ | |
| | HMBS$^{L,S}$ | | NAGLU$^{S}$ | |
| | INPP5B$^{C}$ | | NME7$^{C,S}$ | |
| | KIAA0040$^{E}$ | | NPHP3$^{S}$ | |

$^{E}$ eCAVIAR , $^{C}$ COLOC, $^{L}$ LD, $^{S}$ SMR

**Supplementary Figure 7: Summary of the four different approaches used to perform SMC eQTL and CAD GWAS colocalization. 1)** eQTLs identified in quiescent and proliferative SMCs were colocalized based on linkage disequilibrium of the CAD index SNP and the most significantly associated eQTL SNP (LD). **2)** They were also colocalized using Summary Level Mendelian Randomization (SMR)[49] to test for the pleiotropic association of gene expression in SMCs and CAD. **3)** eQTL and GWAS CAusal Variants Identification in Associated Regions (eCAVIAR)[50] was used to test for causal SNPs between SMC eQTLs and the CAD GWAS. **4)** Bayesian Colocalization Analysis (COLOC)[51] was used to identify the colocalization between SMC eQTLs and the CAD GWAS. The letters in superscript for each gene indicate the approach where the evidence for the colocalization comes from.

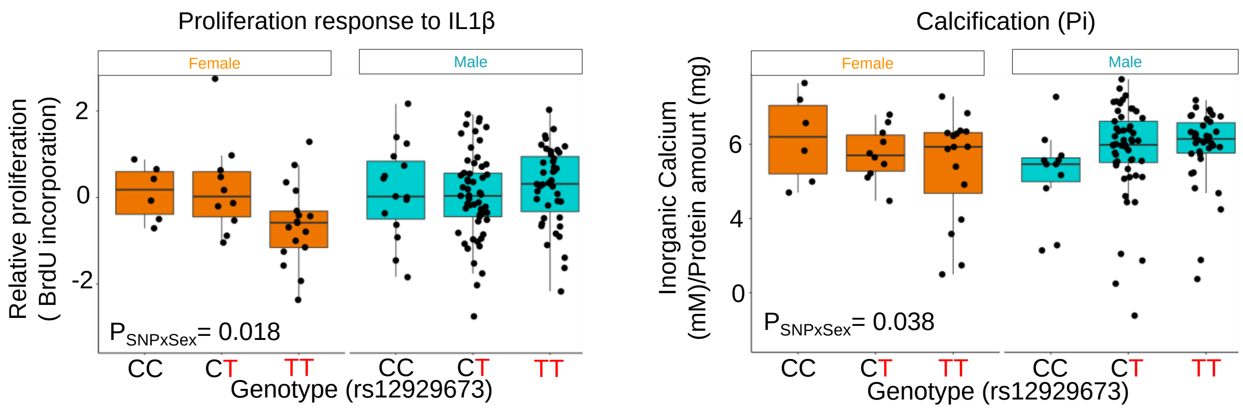


**Supplementary Figure 8: Sex-biased association of the *TERF2IP* locus with SMC proliferation and calcification.** We have previously characterized the same SMC donors used in this study for 12 atherosclerosis-relevant phenotypes[69]. The associations of the rs12929673 with proliferation response to IL-1β and calcification in inorganic phosphate cell culture medium are shown.

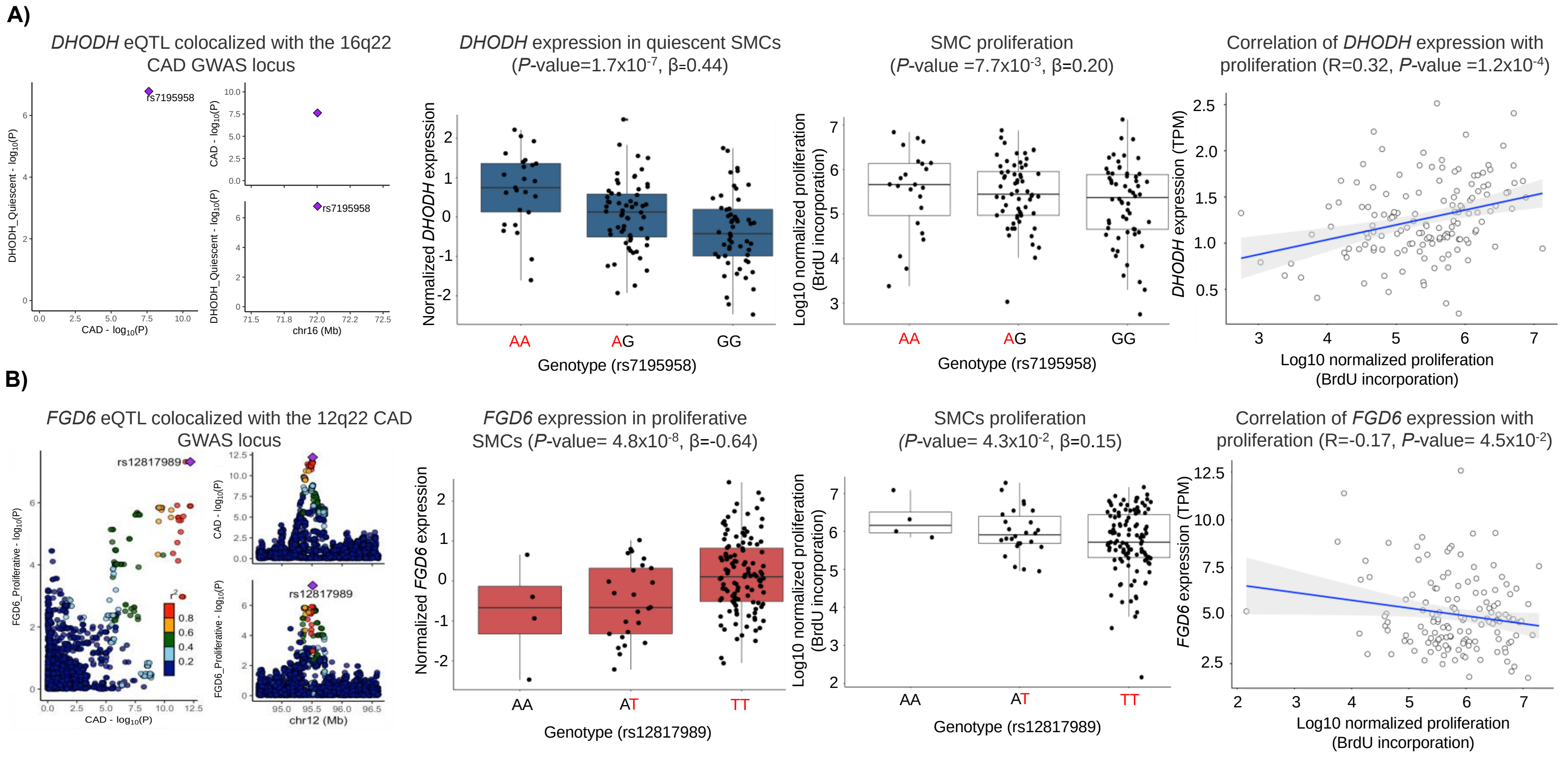


**Supplementary Figure 9: Association of CAD colocalized SMC cis-eQTLs with atherosclerosis-relevant SMC phenotypes. A)** *DHODH cis*-eQTL colocalized with the 16q22 CAD GWAS locus (left), the risk allele, A, of the SNP rs7195958 is associated with higher *DHODH* expression in quiescent SMCs and also SMC proliferation. We observed significant positive correlation between *DHODH* expression and SMC proliferation (right). **B)** *FGD6 cis*-eQTL signal colocalized with the 12q22 CAD GWAS locus (left), the risk allele, T, of the SNP rs12817989 is associated with higher *FGD6* expression in proliferative SMCs and lower proliferation. We observed a significant negative correlation between *FGD6* expression and SMC proliferation (right).

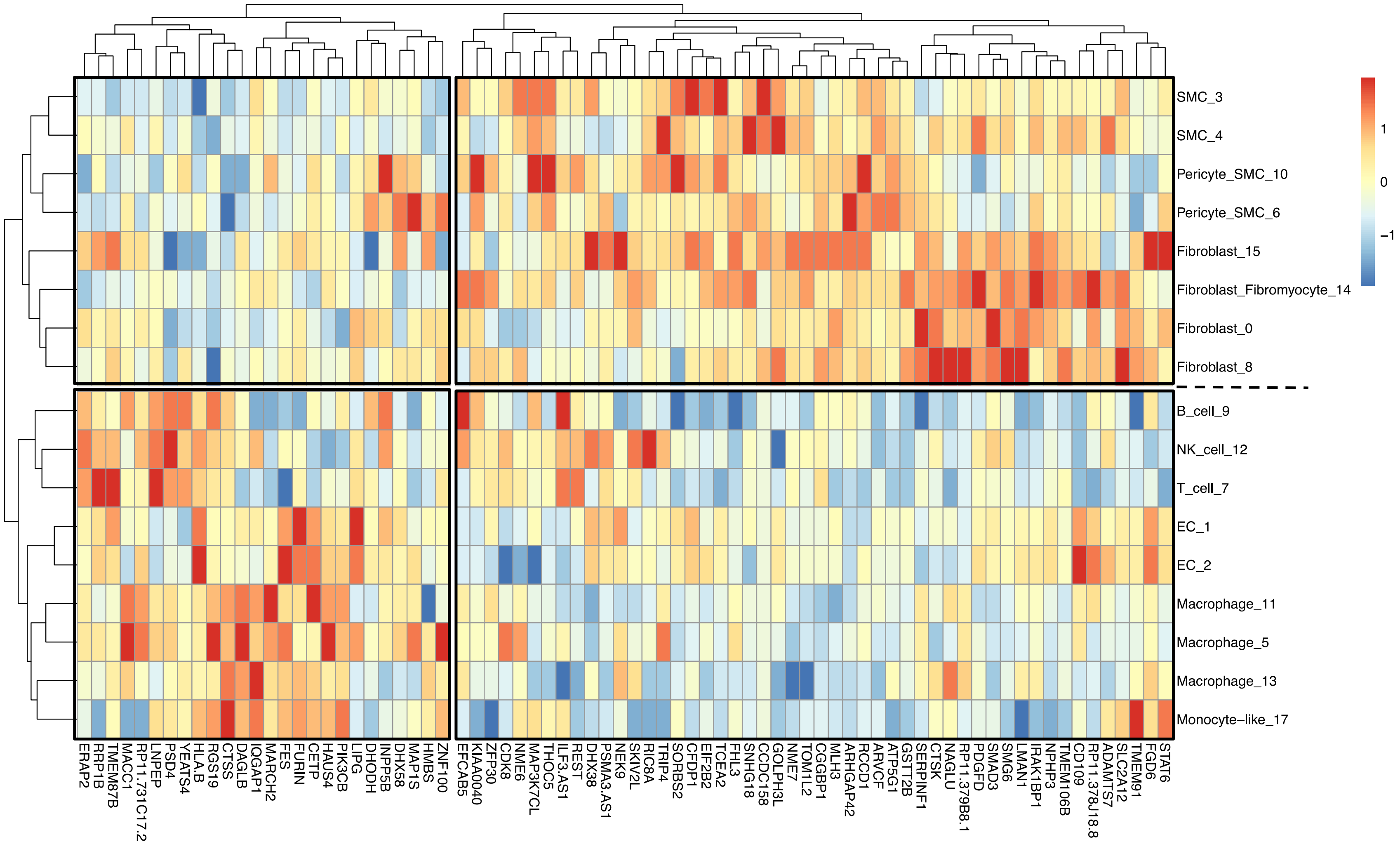


**Supplementary Figure 10: SMC eQTL gene expression in single cells isolated from human coronary atherosclerotic plaques.** The scRNAseq data has been published previously[70]. We were able to assess the expression of 76 of the 84 SMC eQTL genes colocalized with CAD loci. 50 of the 76 genes had higher expression in SMCs, pericytes, and fibroblast compared to endothelial cells, monocytes, macrophages, and other immune cells. The color key of the amount of expression ranging from −2 (blue) to +2 (red) is shown on the right. Red color indicates higher expression, while blue color indicates lower expression.

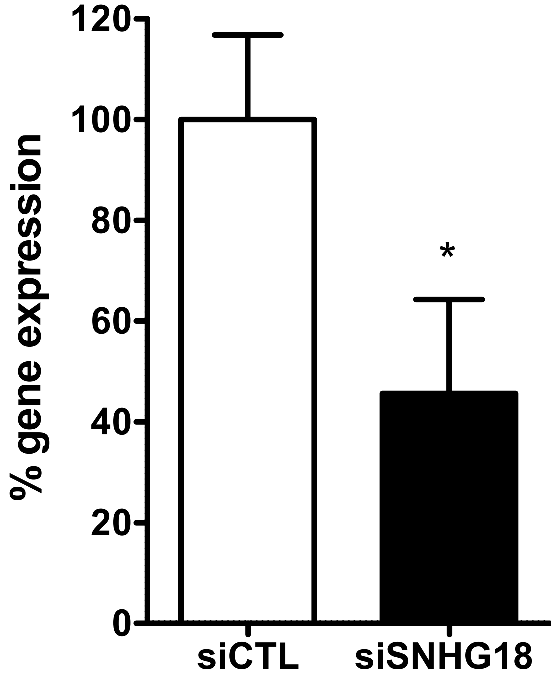


**Supplementary Figure 11: qPCR results after SNHG18 downregulation in SMCs.** SMCs were transfected with control siRNA, and SNHG18 siRNA and qPCR analyses were conducted 48 hours post-transfection.

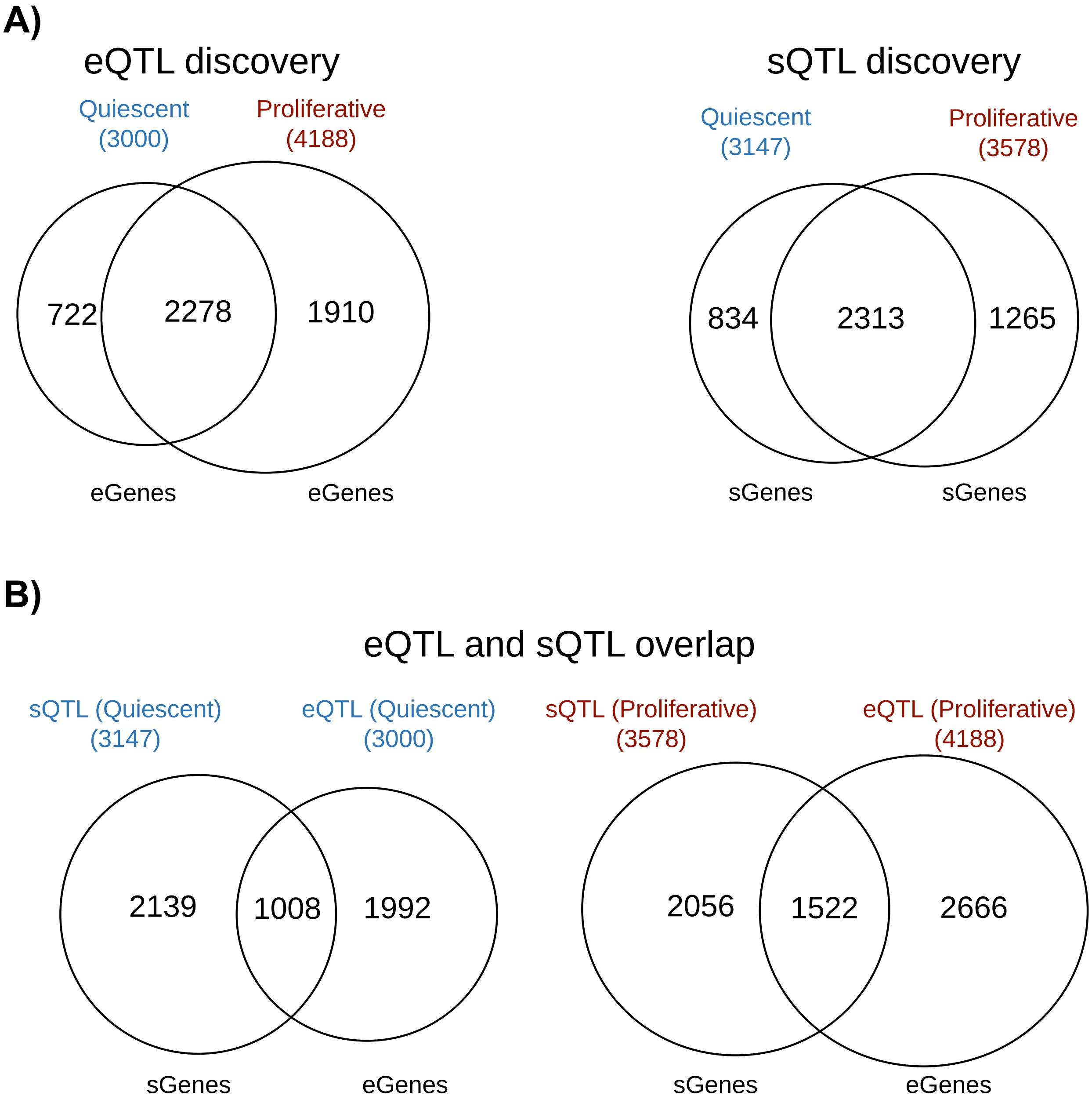


**Supplementary Figure 12: Number of eGenes and sGenes discovered in quiescent and proliferative SMCs. A)** Comparison of the number of eGenes and sGenes discovered in quiescent and proliferative SMCs showed a large overlap between the phenotypes. **B)** Overlap of genes with an eQTL or sQTL showed that genetic regulation of the expression or splicing of mRNAs was largely independent.
